# Recognition of Z-RNA by ADAR1 limits interferon responses

**DOI:** 10.1101/2020.12.04.411793

**Authors:** Qiannan Tang, Rachel E. Rigby, George R. Young, Astrid Korning-Hvidt, Tiong Kit Tan, Anne Bridgeman, Alain R. Townsend, George Kassiotis, Jan Rehwinkel

## Abstract

Nucleic acids are powerful triggers of innate immunity and can adopt the unusual Z-conformation. The p150 isoform of adenosine deaminase acting on RNA 1 (ADAR1) prevents aberrant interferon (IFN) induction and contains a Z-nucleic acid binding (Z*α*) domain. We report that knock-in mice bearing two point mutations in the Z*α* domain of ADAR1, which abolish binding to Z-form nucleic acids, spontaneously induced type I IFNs and IFN-stimulated genes (ISGs) in multiple organs. This included the lung where both stromal and haematopoietic cells displayed ISG induction in *Adar1^mZα/mZα^* mice. Concomitantly, *Adar1^mZα/mZα^* mice showed improved control of influenza A virus. The spontaneous IFN response in *Adar1^mZα/mZα^* mice required MAVS, implicating cytosolic RNA sensing. Finally, analysis of A-to-I changes revealed a specific requirement of ADAR1’s Z*α* domain in editing of a subset of RNAs. In summary, our results reveal that endogenous RNAs in Z-conformation have immunostimulatory potential that is curtailed by ADAR1.

## Introduction

The innate immune system monitors the intra- and extracellular environments for unusual nucleic acids (Bartok and Hartmann, 2020). This process, known as ‘nucleic acid sensing’, detects pathogen invasion and disturbances to homeostasis. It involves a large number of germline encoded nucleic acid sensors. Upon engagement by immunostimulatory DNA or RNA, these sensors signal to initiate a large spectrum of responses, including transcription of the genes encoding type I interferons (IFNs). Type I IFNs – secreted cytokines that act in paracrine and autocrine manner – induce expression of hundreds of IFN-stimulated genes (ISGs). The proteins encoded by ISGs mediate a plethora of functions and include antiviral effectors (Schoggins, 2019). Sustained type I IFN responses can have detrimental effects and cause a range of diseases including the neuroinflammatory Aicardi-Goutières Syndrome (AGS) (Uggenti et al., 2019). It is therefore important to understand the molecular mechanisms that prevent activation of nucleic acid sensors by ‘normal’ DNA and RNA present in healthy cells (Bartok and Hartmann, 2020).

We and others proposed that double-stranded (ds) nucleic acids adopting an unusual conformation known as Z-DNA/RNA activate innate immunity (Maelfait et al., 2017; Sridharan et al., 2017; Zhang et al., 2020b). Z-DNA was initially described by Alexander Rich (Wang et al., 1979). Unlike canonical B-DNA, a right-handed double helix, Z-DNA is a left-handed double helix with a zig-zag-shaped phosphodiester back bone (Wang et al., 1979). dsRNA can also adopt the Z-conformation (Davis et al., 1986; Hall et al., 1984). Biological functions of Z nucleic acids, in particular those of Z-RNA, are incompletely understood (Herbert, 2019). A small number of proteins, all involved in innate immunity, contain Z-DNA/RNA binding domains known as Z*α* domains (Athanasiadis, 2012). These domains specifically bind to and stabilise Z-DNA/RNA, or induce the Z-conformation (Athanasiadis, 2012; Brown et al., 2000; Kim et al., 2018; Schwartz et al., 1999).

There are two mammalian proteins with Z*α* domains: Z-DNA binding protein 1 (ZBP1) and adenosine deaminase acting on RNA 1 (ADAR1, also known as DRADA1). ZBP1 contains two Z*α* domains that have been suggested to recognise viral and endogenous Z-RNAs (Devos et al., 2020; Jiao et al., 2020; Maelfait et al., 2017; Sridharan et al., 2017; Wang et al., 2020; Zhang et al., 2020b). Binding to Z-RNA activates ZBP1 and results in the induction of necroptosis, an inflammatory form of cell death (Maelfait et al., 2020).

ADAR1 has two splice isoforms: ADAR1-p110, which is constitutively expressed and localised in the cell nucleus, and ADAR1-p150 that is IFN-inducible and present in the nucleus and cytosol. Both isoforms contain a C-terminal deaminase domain that converts adenosine to inosine in dsRNA, a process known as A-to-I RNA editing. Conversion of adenosine to inosine in protein-coding sequences can lead to incorporation of non-synonymous amino acids during translation due to base pairing of inosine with cytosine. However, the vast majority of A-to-I editing events occur in non-coding RNAs (Eisenberg and Levanon, 2018; Reich and Bass, 2019). This includes transcripts from repetitive elements (REs), particularly *Alu* elements and short interspersed nuclear elements (SINEs) in human and mouse, respectively. Both ADAR1 isoforms further contain three dsRNA binding domains (dsRBDs) and a so-called Z*β* domain. Z*β* adopts a fold similar to Z*α* domains but does not bind Z-form nucleic acids due to substitutions of key amino acids (Athanasiadis et al., 2005; Kim et al., 2003). ADAR1-p150 has an extended N-terminus harbouring a Z*α* domain (Heraud-Farlow and Walkley, 2020).

ADAR1 deficiency results in profound inflammatory phenotypes. In human, germline *ADAR1* mutations cause AGS (Rice et al., 2012). These mutations predominantly map to the deaminase domain; however, one mutation encoding p.Pro193Ala is found in the Z*α* domain. Pro193 contributes to Z-form nucleic acid binding (Schwartz et al., 1999) and changing it to Ala reduces RNA editing in a reporter assay (Mannion et al., 2014). *Adar1^-/-^* mice, editing-deficient *Adar1^E861A/E861A^* knock-in animals and *Adar1^p150-/p150-^* mice, which only lack ADAR1-p150, all die *in utero* (Hartner et al., 2004; Liddicoat et al., 2015; Wang et al., 2004; Ward et al., 2011). Akin to spontaneous type I IFN induction in AGS patients with *ADAR1* mutation, *Adar1^-/-^* and *Adar1^E861A/E861A^* embryos display profound type I IFN responses prior to death (Hartner et al., 2009; Liddicoat et al., 2015). Multiple nucleic acid sensors mediate the anti-proliferative, cell death and type I IFN phenotypes observed in ADAR1-deficient settings: the oligoadenylate synthetase (OAS)–RNase L system (Li et al., 2017), protein kinase R (PKR) (Chung et al., 2018; Li et al., 2010) and melanoma differentiation-associated protein 5 (MDA5) (Liddicoat et al., 2015; Mannion et al., 2014; Pestal et al., 2015). Upon activation by dsRNA, OAS proteins synthesise 2’-5’ oligoadenylate, a second messenger that in turn activates RNase L, resulting in widespread RNA degradation. PKR also detects dsRNA and represses protein translation. Both effects have been proposed to explain lethality of ADAR1-deficient cells (Chung et al., 2018; Li et al., 2017). Induction of type I IFNs in ADAR1-deficient mice and human cells is mediated by the RNA sensor MDA5, that signals via its adaptor mitochondrial antiviral-signalling protein (MAVS) (Bajad et al., 2020; Chung et al., 2018; Guallar et al., 2020; Liddicoat et al., 2015; Mannion et al., 2014; Pestal et al., 2015).

These observations suggest a model in which endogenous dsRNAs are stabilised in ADAR1-deficient cells due to the absence of RNA editing and are then recognised by RNA sensors (Dias Junior et al., 2019; Eisenberg and Levanon, 2018). Indeed, some ADAR1 substrates such as transcripts from *Alu* elements base-pair to form duplex structures, which are predicted to be destabilised by inosine : uracil mismatches introduced by RNA editing (Ahmad et al., 2018; Chung et al., 2018; Pfaller et al., 2018; Song et al., 2020). This concept has been tested experimentally in an RNase protection assay: transcripts spanning *Alu* elements in inverted orientation are protected by recombinant MDA5 protein in RNA samples extracted from ADAR1-deficient cells (Ahmad et al., 2018; Mehdipour et al., 2020).

How the different nucleic acid binding domains in ADAR1 select and recruit RNA substrates for subsequent editing is unknown. In light of (i) emerging roles of Z-form nucleic acids in innate immunity (Kesavardhana and Kanneganti, 2020), (ii) the MAVS-dependent phenotype of *Adar1^p150-/p150-^* mice (Pestal et al., 2015), and (iii) natural occurrence of the p.Pro193Ala mutation in AGS patients (Rice et al., 2012), we interrogated Z-form nucleic acid binding by the Z*α* domain in ADAR1-p150. Here, we report the generation of knock-in mice bearing two missense mutations in the Z*α* domain, which prevent nucleic acid binding. Although these *Adar1^mZα/mZα^* mice were developmentally normal and fertile, they displayed spontaneous induction of type I IFNs and ISGs in multiple organs and cell types. This phenotype was dependent on MAVS and conferred partial protection against influenza A virus (IAV) infection. Analysis of sequencing data revealed that ∼8% of RNA editing events in wild-type (WT) cells required a functional ADAR1-p150 Z*α* domain. Taken together, our study delineates the function of the Z*α* domain in ADAR1-p150 *in vivo* and suggests that recognition of Z-form RNA by ADAR1 contributes to the suppression of IFN responses.

## Results

### Generation of Z*α* domain mutated mice

To study the role of Z-form nucleic acid binding to the Z*α* domain in ADAR1-p150 in an *in vivo* setting, we generated knock-in mice bearing two missense mutations: p.Asn175Ala and p.Tyr179Ala. These residues are conserved and are homologous to Asn173 and Tyr177 in human ADAR1; they were chosen because of their essential role in Z-form nucleic acid binding (Feng et al., 2011; Li et al., 2009; Schade et al., 1999; Schwartz et al., 1999). Given the embryonic lethality of *Adar1^p150-/p150-^* mice (Ward et al., 2011), we opted for a conditional knock-in strategy (Figure S1A). The Z*α* domain is encoded by exon 2 of the *Adar1* gene. In brief, we introduced in inverted orientation into the intron between exons 2 and 3 a mutated copy of exon 2 (designated 2*) containing four nucleotide substitutions, changing both Asn175 and Tyr179 to Ala. We flanked exons 2 and 2* with LoxP and Lox2272 sites such that Cre-mediated recombination removes exon 2 and flips exon 2* into forward orientation (Figure S1A). We designated the conditional allele ‘*fl-mZα*’ and the recombined allele expressing mutant Adar1 ‘*mZα*’. To determine the impact of the Z*α* domain mutations when present in all cells and tissues, we crossed *Adar1^+/fl-mZα^* mice with a line expressing Cre recombinase under control of the ubiquitously active *Pgk* promoter. The resulting *Adar1^+/mZα^* mice were intercrossed to generate homozygous animals. We validated the presence of the mutations by sequencing and found that *Adar1^mZα/mZα^* animals were born at expected mendelian ratios (Figure S1B and S1C). Furthermore, they developed normally and were fertile. The mutations introduced into the Z*α* domain did not alter the expression of the two isoforms of ADAR1 in bone marrow-derived myeloid cells (BMMCs) (Figure S1D). Taken together, Z-form nucleic acid binding by ADAR1-p150 is not essential for survival at whole organism level.

### Mutation of the Z*α* domain in ADAR1-p150 triggers a multi-organ type I IFN response

We tested whether *Adar1^mZα/mZα^* mice display spontaneous activation of type I IFNs as was reported in ADAR1-deficient settings. We collected lung, liver and spleen tissues and extracted RNA for RT-qPCR analysis. Transcript levels of *Ifnb1* (encoding IFN*β*) were elevated in lung RNA samples from *Adar1^mZα/mZα^* mice (Figure 1A). Moreover, the ISG transcripts *Ifit1* and *Ifi44* were expressed at higher levels in all three organs from Z*α* domain mutated animals (Figure 1A). This gene expression signature was type I IFN-specific: tissues from *Adar1^mZα/mZα^* mice contained comparable mRNA levels of *Ifng*, *Tnfa* and *Ιl1b* (encoding IFN*γ*, TNF*α* and pro-IL-1*β*), with only a minor increase of *Tnfa* mRNA in lung (Figure 1A). We validated the ISG signature at protein level by analysing expression of interferon stimulated gene 15 (ISG15) in whole lung lysates. ISG15 is a ubiquitin-like modifier that is induced by IFN and is conjugated to target proteins in a process called ISGylation (Perng and Lenschow, 2018). Western blot showed that lung lysates from *Adar1^mZα/mZα^* mice contained increased amounts of monomeric ISG15 as well as increased levels of ISGylated proteins, visible as a high-molecular weight smear (Figure 1B).

**Figure 1.**
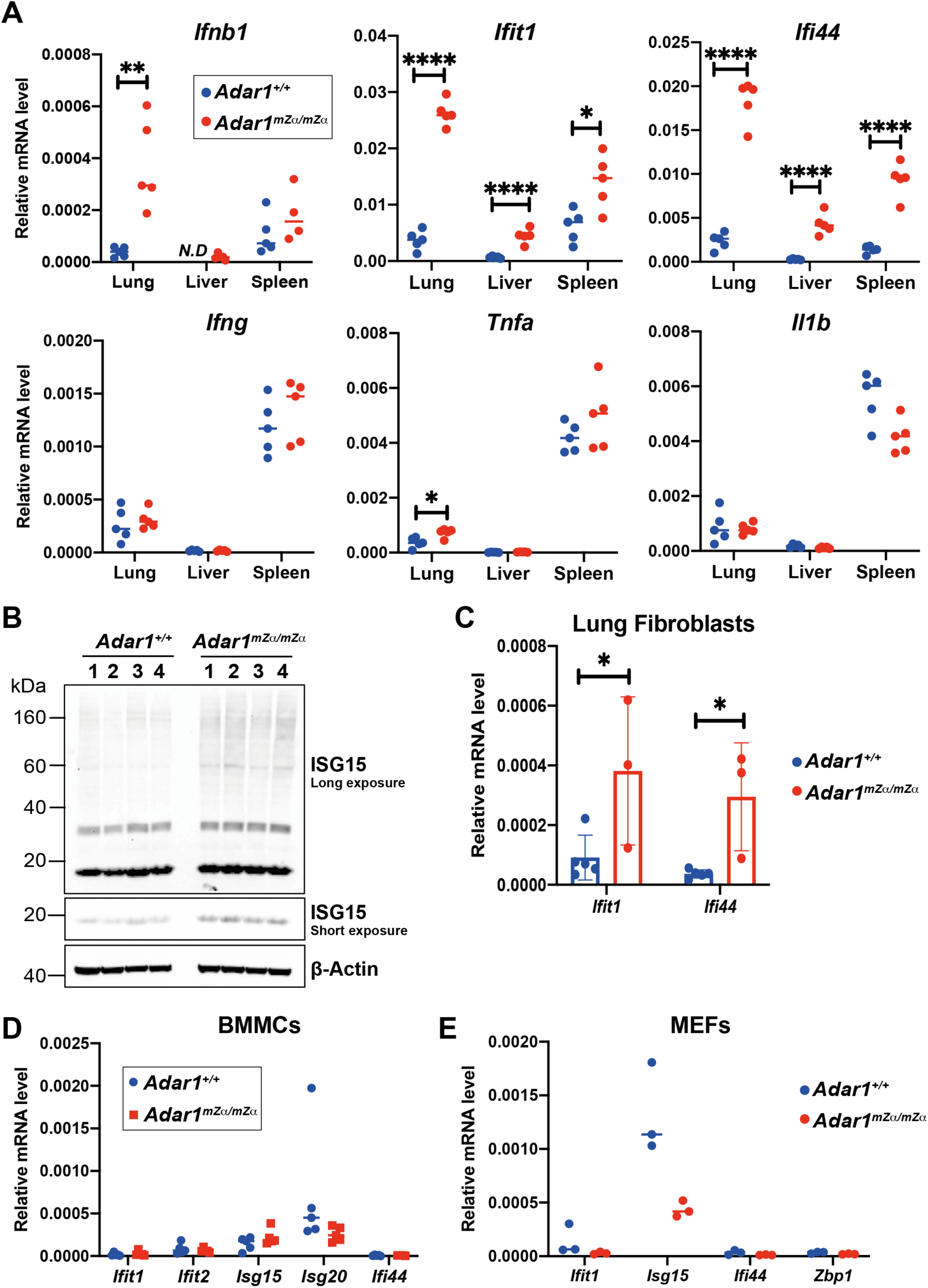
Mutation of ADAR1-p150’s Z*α* domain triggers spontaneous type I IFN responses in multiple organs. **A.** Levels of the indicated mRNAs were analysed by RT-qPCR in RNA samples extracted from tissues of WT and *Adar1^mZα/mZα^* animals and are shown relative to *Gapdh*. Each dot represents an individual mouse. *N.D*, not detectable. **B.** Protein extracts from whole lungs from animals of the indicated genotypes were used for western blot with an *α*-ISG15 antibody. *β*-Actin served as a loading control. Each lane represents a sample from an individual mouse. **C-E.** mRNA levels of the indicated ISGs were analysed by RT-qPCR from cultured lung fibroblasts (C), BMMCs (D), and MEFs (E) of the indicated genotypes and are shown relative to *Gapdh*. Each dot represents cells derived from an individual mouse. Pooled data from biological replicates are shown with mean (A, D, E) or mean ± SD (C) and were analysed by unpaired t test (****p<0.0001, **p<0.01, *p<0.05). See also Figure S1.

We next analysed different types of cultured primary cells from WT and *Adar1^mZα/mZα^* mice. We observed heightened ISG expression in Z*α* domain mutated lung fibroblasts (Figure 1C). In contrast, comparable levels of ISG transcripts were found in WT and *Adar1^mZα/mZα^* BMMCs, as well as in mouse embryonic fibroblasts (MEFs) (Figure 1D, E). This indicates that ISG induction may be cell-type specific.

Among the organs we analysed, lung exhibited the most profound ISG signature, and spontaneous ISG induction was also observed in cultured lung fibroblasts (Figure 1A, D). We therefore focused on the lung for the next set of experiments. We used RNA extracted from lung for RNA sequencing to obtain a global view of gene expression in *Adar1^mZα/mZα^* mice. We found that 99 protein coding genes were differentially expressed by at least 2-fold in mutant lungs, including 89 upregulated and 10 downregulated transcripts (Figure 2A). 40% of the induced mRNAs were encoded by ISGs; this included well-known factors such as *Irf7*, *Cxcl10*, *Zbp1* and *Usp18* (Figure 2A). We used gene ontology (GO) analysis of biological processes to further classify upregulated genes. GO terms related to type I IFNs and anti-viral defence were enriched amongst genes induced in *Adar1^mZα/mZα^* lungs (Figure 2B). The most highly enriched GO category, regulation of ribonuclease activity, included many *Oas* transcripts, which are known to be IFN-inducible (Schoggins, 2019).

**Figure 2.**
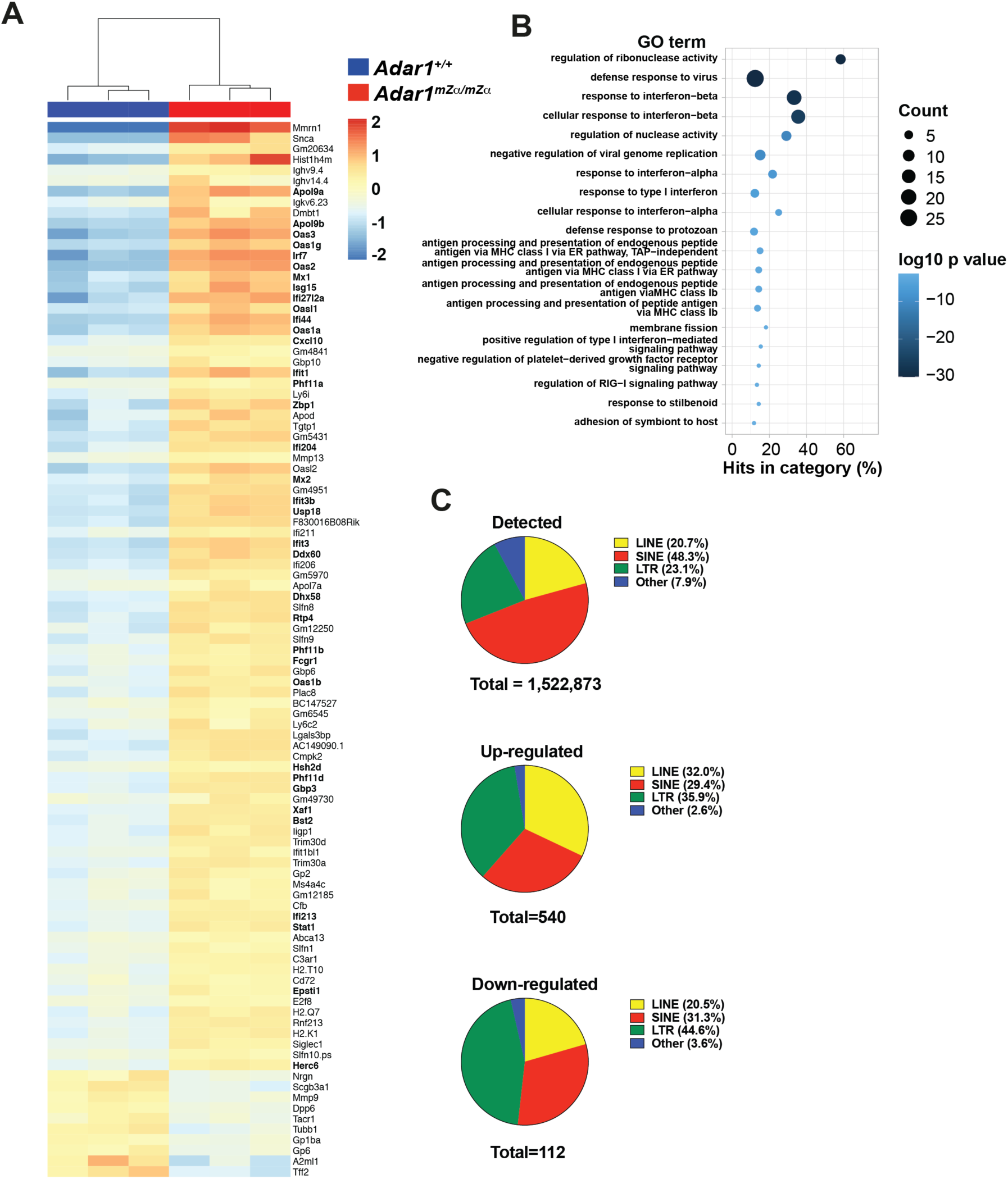
*Adar1^mZα/mZα^* lungs display a type I IFN gene signature. Total RNA was extracted from lungs of three WT and three *Adar1^mZα/mZα^* mice. Ribosomal RNAs were depleted before random-primed library preparation and RNA sequencing. About 100 million reads were obtained per sample. **A.** Differentially expressed genes were defined as displaying a fold change of ≥2 with an adjusted p-value <0.01. The 89 upregulated and 10 downregulated genes were ordered by decreasing fold change and the data were clustered by sample. ISGs are indicated in bold. **B.** GO analysis of up-regulated genes. The top 20 GO terms (biological processes), ranked and ordered by p-value, are shown. Diameters indicate the number of induced genes assigned to the GO term and colours show the p-value. **C.** Detected and differentially expressed REs were assigned to the indicated classes and are shown as pie charts. Differentially expressed REs were identified as having a minimum fold change of 2 and an adjusted p value of less than 0.01. See also Table S1.

We also analysed REs in our RNA sequencing data and found 540 induced and 112 repressed REs (Figure 2C and Table S1). SINEs were underrepresented amongst differentially expressed REs, whilst long terminal repeat (LTR) elements were enriched (Figure 2C). Dysregulation of REs has been reported in settings of inflammation and can be driven by IFNs (Chuong et al., 2016). Taken together, these data show that the lungs of Z*α* domain mutated animals displayed a type I IFN-driven gene signature.

### Stromal and haematopoietic cells contribute to the ISG signature in the lungs of *Adar1^mZα/mZα^* mice

To identify the type of cell(s) that display the ISG signature in the lungs of *Adar1^mZα/mZα^* mice, we used magnetic-activated cell sorting (MACS) to isolate haematopoietic cells, marked by CD45 expression, and stromal cells lacking this marker. There was no difference in the proportions of haematopoietic and stromal cells between WT and *Adar1^mZα/mZα^* lungs (Figure 3A). We confirmed the purity of MACS-enriched haematopoietic and stromal cells to be >97% (Figure S2). We then extracted RNA for RT-qPCR analysis of ISGs and found that both lung haematopoietic and stromal cells displayed ISG induction (Figure 3B). For the three ISGs tested, fold inductions in *Adar1^mZα/mZα^* samples were similar in CD45+ and CD45-cells (Figure 3B). However, it is noteworthy that baseline ISG levels appeared to be lower in haematopoietic cells compared to stromal cells (Figure 3B).

**Figure 3.**
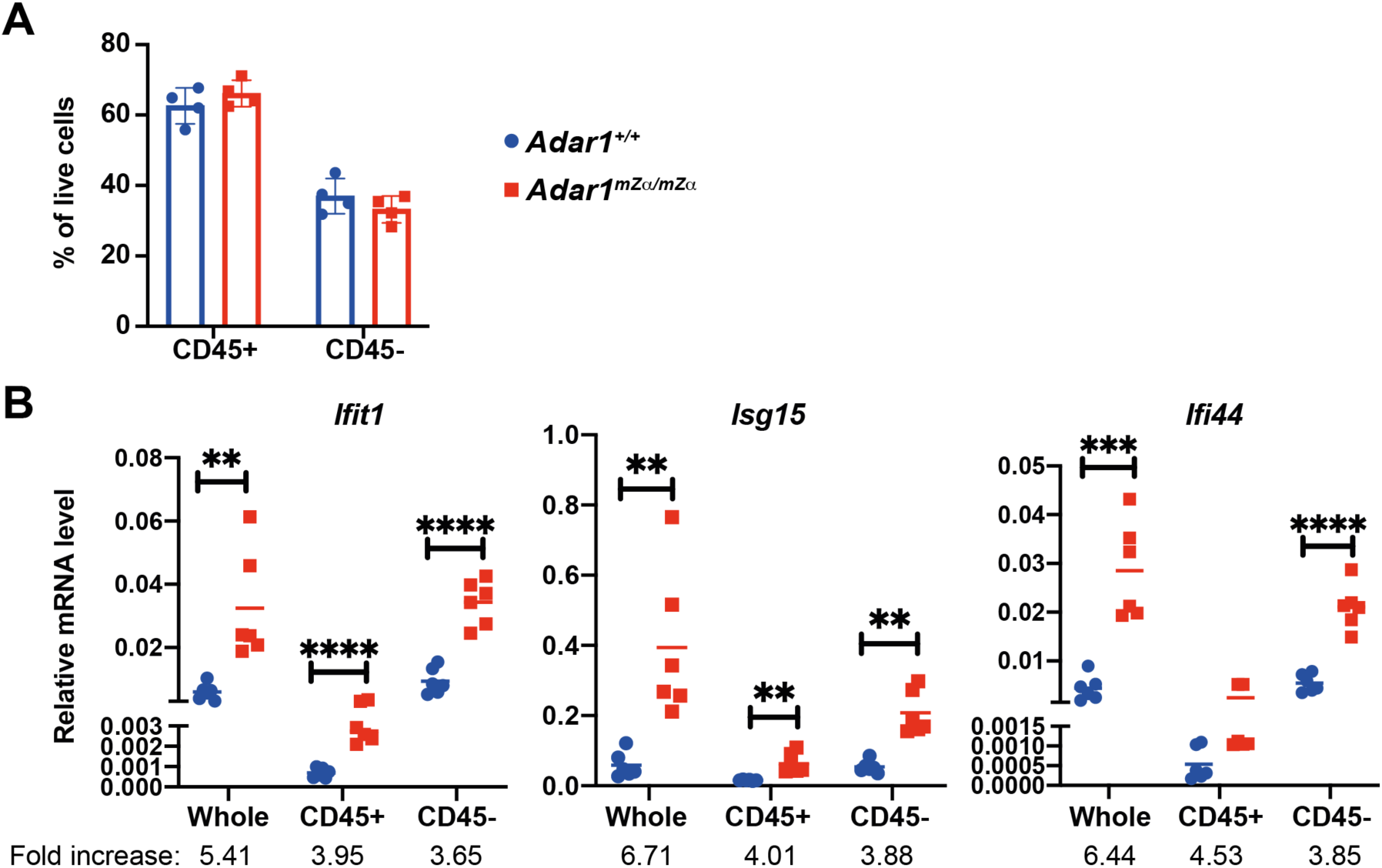
Stromal and haematopoietic cells upregulate ISGs in *Adar1^mZα/mZα^* lungs. **A.** The proportion of haematopoietic (CD45+) and stromal (CD45-) cells in WT and *Adar1^mZα/mZα^* lungs is shown. **B.** mRNA levels of the indicated ISGs were analysed by RT-qPCR using RNA extracted from whole lung, or from CD45+ or CD45-cells, and are shown relative to *Actb*. Fold increases relative to WT samples were calculated. Data points represent individual animals. In (A), data from a representative experiment are shown with mean ± SD. In (B), pooled data from two independent experiments including a total of six animals are shown with mean and were analysed by unpaired t test (****p < 0.0001, ***p<0.001, **p<0.01). See also Figure S2.

Next, we analysed ISG expression in different types of lung haematopoietic and stromal cells. Using two staining panels and fluorescence-activated cell sorting (FACS), we obtained eight different populations of CD45+ cells, including B cells, T cells, dendritic cells (DCs), monocytes, macrophages, NK cells, neutrophils and eosinophils (Figure S3). There were no differences between WT and *Adar1^mZα/mZα^* mice in the proportions of these cell populations, with the exception of DCs, which were less abundant in the mutant lungs (Figure S3A). *Ifit1*, *Ifit2* and *Isg15* were significantly induced in neutrophils from the lungs of *Adar1^mZα/mZα^* mice (Figure 4A, B). However, these ISGs showed no, or limited and statistically insignificant, upregulation in other types of CD45+ cells (Figure 4A, B). In contrast, the ISGs *Ifi44* and *Oas1a* were upregulated in multiple haematopoietic cell types (Figure 4A, B). Transcripts encoding type I IFNs were undetectable by RT-qPCR in all samples analysed (data not shown). Several ISGs, including *Ifit1*, *Ifit2* and *Isg15*, are not only induced by IFNAR signalling but can also be upregulated by IRF3/7, independent of type I IFN-initiated JAK-STAT activation (Ashley et al., 2019; DeFilippis et al., 2006; Goubau et al., 2009; Grandvaux et al., 2002; Lazear et al., 2013). Induction of this ISG-subset can therefore occur in a cell-autonomous manner, for example when aberrant nucleic acids are sensed by pattern recognition receptors (PRRs) that signal via IRF3/7. Other ISGs, including *Ifi44* and *Oas1a*, are only induced via IFNAR signalling (Lazear et al., 2013). Therefore, our observation that *Ifit1*, *Ifit2* and *Isg15* were primarily induced in neutrophils, while *Ifi44* and *Oas1a* were upregulated more broadly (Figure 4A, B), suggests that *Adar1^mZα/mZα^* neutrophils may autonomously activate IRF3. We further speculate that the ISG signature in other haematopoietic cells such as B cells may be due to paracrine type I IFN signalling via IFNAR.

**Figure 4.**
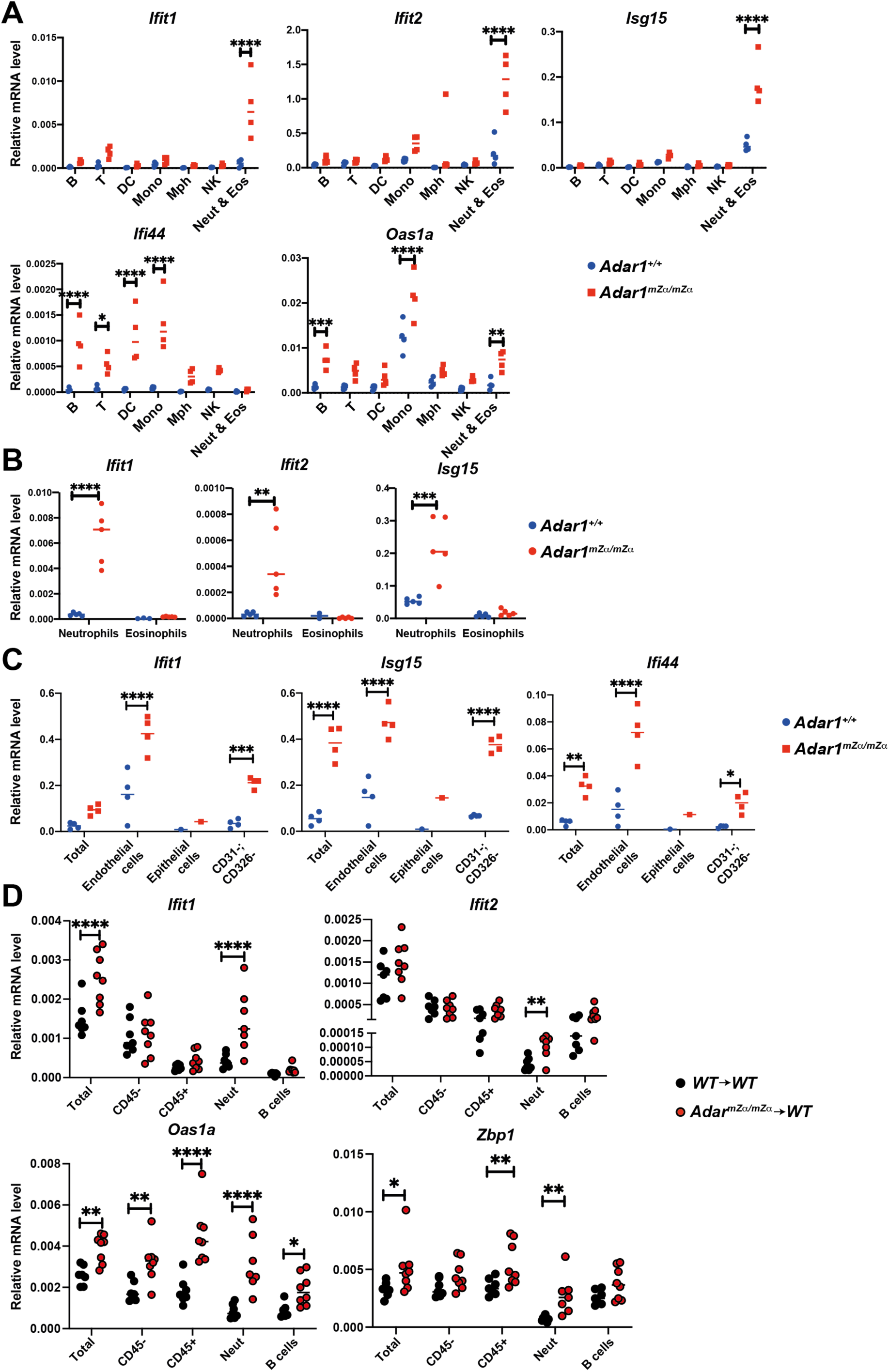
Multiple haematopoietic and non-haematopoietic cell types display ISGs upregulation in *Adar1^mZα/mZα^* lungs. **A-C.** mRNA levels of the indicated ISGs were analysed by RT-qPCR using RNA extracted from cell populations sorted from lungs of WT and *Adar1^mZα/mZα^* mice and are shown relative to *Actb*. B, B cells; T, T cells; DC, dendritic cells; Mono, monocytes; Mph, macrophages; NK, natural killer cells; Neut, neutrophils; Eos, eosinophils; Total, whole lung. **D.** ISG mRNA levels were analysed as in (A-C) in cell populations sorted from lungs of BM chimeric mice and are shown relative to *Gapdh*. Each data point represents an individual mouse. Due to the small number of epithelial cells recovered, samples were pooled from multiple mice before RNA extraction (C). Pooled data from two (A,B) or three (D) independent experiments are shown with mean (****p < 0.0001, ***p<0.001, **p<0.01, *p<0.05, unpaired t test). See also Figures S3, S4 and S5.

We also analysed the lung stromal compartment and isolated by FACS endothelial and epithelial cells using CD31 and CD326 as markers, respectively (Figure S4). We further isolated CD45-cells lacking these markers; this mixed population likely included fibroblasts and other cell types. We observed induction of *Ifit1*, *Isg15* and *Ifi44* in *Adar1^mZα/mZα^* cells of all three populations analysed (Figure 4C). It is therefore likely that, in addition to neutrophils, multiple non-haematopoietic cells initiate type I IFN production in the lungs of *Adar1^mZα/mZα^* mice.

### Haematopoietic cells are sufficient to induce ISGs in *Adar1^mZα/mZα^* mice

To further dissect the cellular requirements for ISG induction in *Adar1^mZα/mZα^* mice, we generated bone marrow (BM) chimeric animals. Lethally irradiated *Cd45.1* WT recipients were reconstituted with *Cd45.2* BM from WT or *Adar1^mZα/mZα^* animals (Figure S5A). We found reconstitution levels of the haematopoietic compartment to be about 90% by analysing cell surface levels of CD45.1 and CD45.2 on white blood cells from recipient mice (Figure S5B). Similar levels of reconstitution were observed in the lung (Figure S5C). We first obtained RNA from whole lung tissue for RT-qPCR analysis. The ISGs *Ifit1*, *Oas1a* and *Zbp1* were upregulated in *Adar1^mZα/mZα^*→WT chimeras compared to WT→WT chimeras (Figure 4D, “Total”). This indicates that *Adar1^mZα/mZα^* BM-derived cells triggered an IFN response in WT animals. We also isolated different cell populations from the lungs of chimeric animals by FACS (see Figure S3 for gating), extracted RNA and performed RT-qPCR. *Ifit1* and *Ifit2* mRNA levels were significantly induced in neutrophils from *Adar1^mZα/mZα^*→WT chimeras, but not in B cells and CD45-cells (Figure 4D). In contrast, *Oas1a* and/or *Zbp1* were upregulated in multiple cell populations including CD45-cells, neutrophils and B cells (Figure 4D). These observations are consistent with the results shown in Figure 4A and 4B and indicate that neutrophils may initiate type I IFN production that then results in ISGs induction in other cell types. Taken together, *Adar1^mZα/mZα^* haematopoietic cells were sufficient to induce ISGs in the lungs of WT mice.

### IAV replication is inhibited in *Adar1^mZα/mZα^* mice

In light of the spontaneous type I IFN response in *Adar1^mZα/mZα^* mice, we next asked whether these Z*α* domain mutant mice were protected against viral infection. ISGs upregulated in the lungs of *Adar1^mZα/mZα^* mice included factors such as *Ifit1* and *Zbp1* that are known to control IAV (Pichlmair et al., 2011; Zhang et al., 2020b) (Figure 2A). We intranasally infected WT and *Adar1^mZα/mZα^* mice with the H3N2 strain of recombinant IAV (A-X31; A/HongKong/1/1968) using a dose that caused moderate weight loss. WT animals lost about 10% body weight from day 3 until day 7 after infection, and subsequently recovered their weight (Figure 5A). In contrast, *Adar1^mZα/mZα^* mice gained weight until day 4 after infection (Figure 5A). This was followed by weight loss on days 5-7, which slightly exceeded weight loss in WT mice, and recovery from day 8 onwards. These observations suggest that *Adar1^mZα/mZα^* mice were protected against IAV at an early stage of the infection.

**Figure 5.**
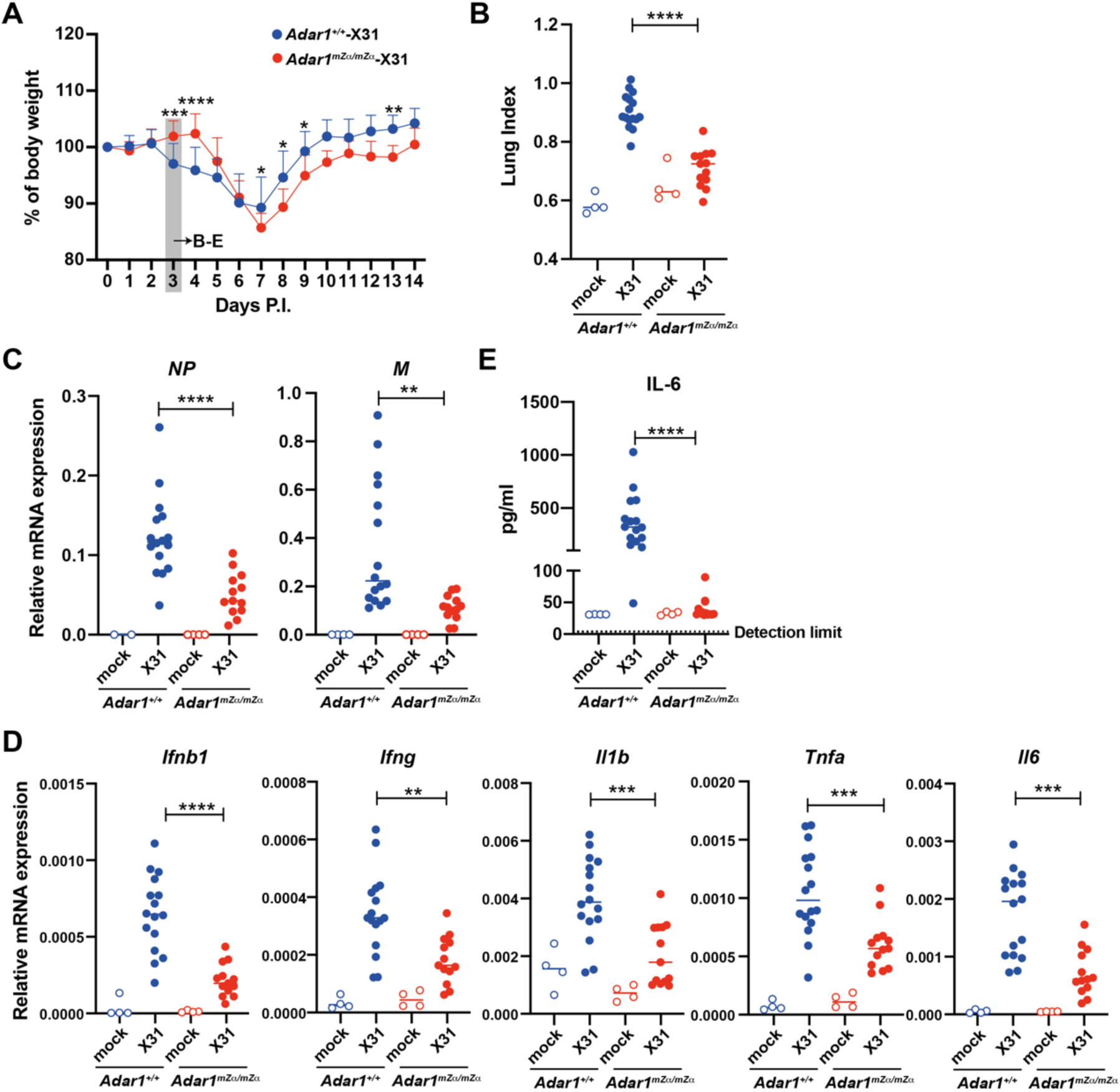
*Adar1^mZα/mZα^* mice are protected from early IAV infection. **A.** WT or *Adar1^mZα/mZα^* mice were infected intranasally with 0.04 HAU of IAV strain A/X31. Body weight was monitored daily and is shown as a percentage of starting body weight. **B-E.** WT or *Adar1^mZα/mZα^* mice were infected as in (A) or mock infected using viral growth medium. On day 3 post infection, lungs and sera were collected. **B.** A ‘lung index’ was calculated (lung weight/body weight x100). **C.** Levels of the viral *NP* and *M* transcripts were analysed by RT-qPCR in RNA samples extracted from total lung. Data are shown relative to *Actb* (*NP*) or *Gapdh* (*M*). **D.** Levels of the indicated mRNAs were determined as in (C). **E.** Serum IL-6 concentrations were analysed by ELISA. In (A), data from three independent experiments including a total of 15 mice per genotype were pooled (mean + SD; ****p<0.0001, ***p<0.001, **p<0.01, *p<0.05, mixed-effects analysis). In (B-E), pooled data from two independent experiments (mock infected: n=4 mice per genotype; X31-infected: n=16 WT and n=13 *Adar1^mZα/mZα^* mice) are shown. Each dot represents an individual mouse and the mean is indicated (****p < 0.0001, ***p<0.001, **p<0.01, unpaired t test).

We further characterised early IAV infection in *Adar1^mZα/mZα^* mice at day 3 after inoculation. We calculated a ‘lung index’ (lung weight/body weight x100; Luo et al., 2012) and found this marker of pathology and inflammation to be significantly higher in infected WT mice compared to *Adar1^mZα/mZα^* animals (Figure 5B). Next, we analysed viral loads by determining the levels of the viral *NP* and *M* transcripts by RT-qPCR. Compared to infected WT lungs, levels of these viral RNAs were reduced in infected *Adar1^mZα/mZα^* lungs (Figure 5C). Concomitantly, mRNA levels of *Ifnb1*, *Ifng*, *Ιl1b*, *Tnfa* and *Il6*, as well as IL6 protein, were strongly induced in infected WT lungs, and these effects were curtailed in *Adar1^mZα/mZα^* lungs (Figure 5D, E). In sum, IAV replication and virus-induced inflammation were reduced in *Adar1^mZα/mZα^* animals.

### ISG induction in *Adar1^mZα/mZα^* mice is MAVS-dependent

We next investigated which nucleic acid sensing pathway triggered spontaneous ISG expression in *Adar1^mZα/mZα^* mice. Since ADAR1-deficiency results in activation of the MDA5-MAVS pathway (Liddicoat et al., 2015; Mannion et al., 2014; Pestal et al., 2015), we hypothesised that the ISG signature in *Adar1^mZα/mZα^* mice was driven by MAVS. To test this, we crossed *Adar1* mutant mice with MAVS knock-out animals to generate *Adar1^mZα/mZα^*; *Mavs^-/-^* mice. Loss of MAVS prevented the ISG induction observed in *Adar1^mZα/mZα^* lungs, livers and spleens (Figure 6A). Because of the neuropathology observed in patients with AGS, we also analysed brain tissue. *Adar1^mZα/mZα^* mice showed elevated *Ifnb1* expression and a pronounced ISG signature in the brain (Figure 6B). Akin to the situation in other tissues, these effects were fully MAVS-dependent (Figure 6B). Taken together, the Z*α* domain of ADAR1-p150 was involved in preventing MAVS-mediated IFN induction, implicating a role in limiting activation of MDA5 or RIG-I.

**Figure 6.**
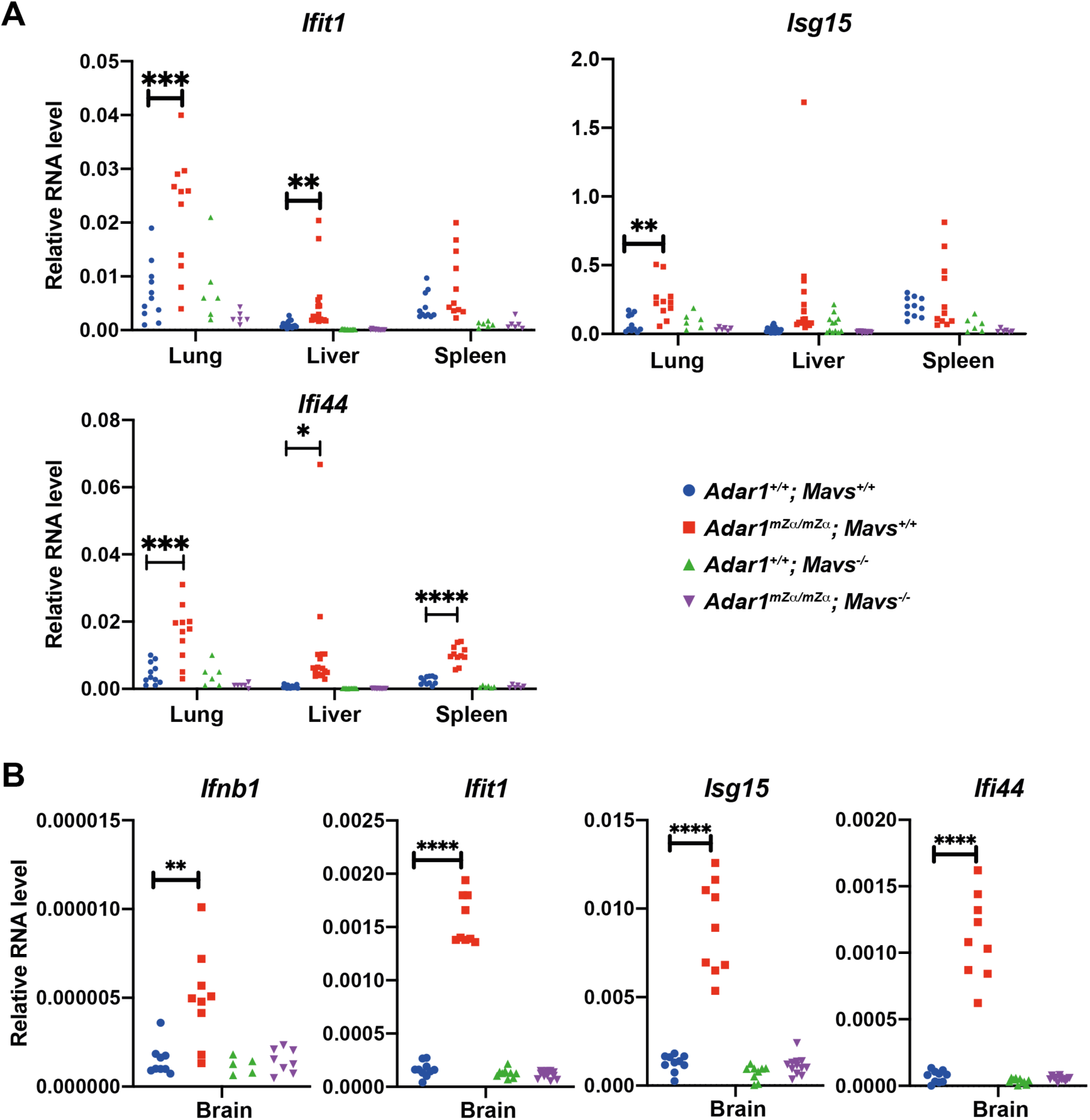
ISG induction in *Adar1^mZα/mZα^* mice is MAVS-dependent. Levels of the indicated mRNAs were analysed by RT-qPCR in RNA samples extracted from tissues of WT and *Adar1^mZα/mZα^* animals that were either MAVS-sufficient or -deficient. Data are shown relative to *Gapdh*. Each dot represents an individual mouse. Pooled data from biological replicates are shown (****p < 0.0001, ***p<0.001, **p<0.01, *p<0.05, unpaired t test).

### The Z*α* domain of ADAR1-p150 is required for editing of a subset of RNAs

Human ADAR1-p150 bearing the p.Pro193Ala mutation shows reduced editing activity in a reporter assay (Mannion et al., 2014). To identify natural RNA substrates edited by ADAR1-p150 in a Z*α* domain-dependent manner, we analysed our RNA sequencing data from WT and *Adar1^mZα/mZα^* lungs for A→G transitions. These are indicative of A-to-I RNA editing as inosine pairs with cytosine during reverse transcription. The most sensitive and specific method for annotating this mutational profile compares the fit of Dirichlet models of observed base frequencies between test and control samples (Piechotta et al., 2017). We extended this methodology to allow comparisons of biological replicates, to incorporate the orientation of the originating RNA (revealed by our stranded RNAseq protocol), to include fine-grained filtering of potential editing sites, and to add a ‘detection’ step, whereby observed base frequencies at potential editing sites are additionally compared to a model of the per-base error rate obtained *de novo* from the dataset.

Considered individually, for each of the three WT and three *Adar1^mZα/mZα^* lung samples analysed, we detected ∼35,000-40,000 editing sites (Figure 7A). Considered as biological replicates, increasing statistical robustness, 40,342 and 46,164 sites were identified that were shared amongst all three WT or *Adar1^mZα/mZα^* lung samples, respectively. These ‘detected’ sites had a median editing level of ∼10% in both WT and *Adar1^mZα/mZα^* samples (Figure 7B). This shows that there was no global defect in RNA editing in *Adar1^mZα/mZα^* mice.

**Figure 7.**
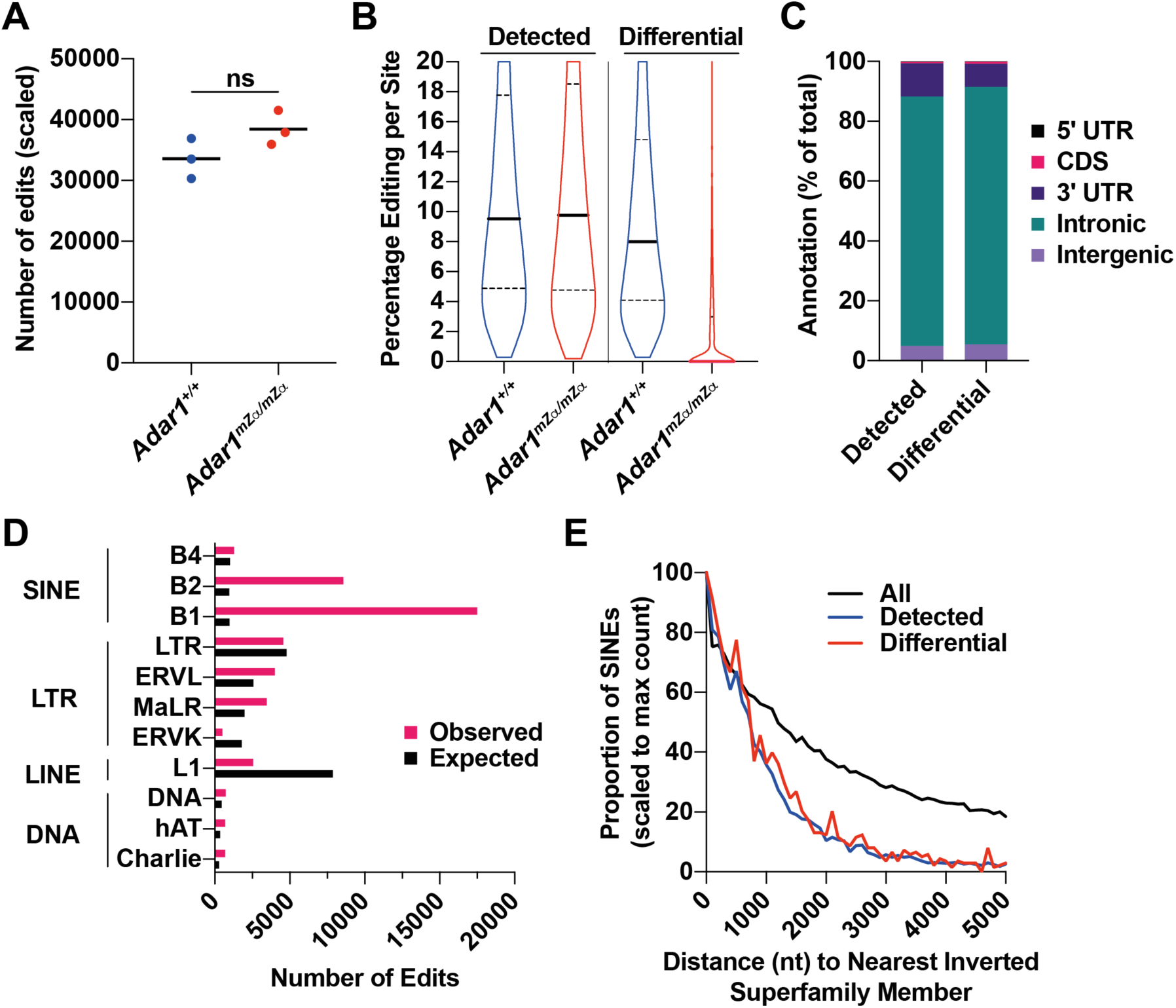
ADAR1-p150’s Z*α* domain is required for editing of a subset of RNAs. **A.** Editing sites were mapped in RNA sequencing reads from three WT and three *Adar1^mZα/mZα^* lung samples (Z>2.58). The numbers of edited sites were scaled to the total number of reads per sample. Each data point corresponds to an animal and the mean is shown (ns, not significant; unpaired t test). **B.** Editing frequencies for sites detectable in all three WT or *Adar1^mZα/mZα^* samples (left; Z>2.58) and for differentially edited sites (right; Z>2.58, >2-fold) are shown as violin plots. Solid horizontal lines show the median and dotted lines indicate quartiles. **C.** Editing sites detected in WT samples and differentially edited sites were matched to annotated genomic features. The percentage of sites is shown for each category. **D.** The number of expected and observed editing sites in WT samples are shown for families of REs for which either value exceeded 500. Please see text for details. **E.** The distances of SINEs to their nearest inverted super-family member were determined for all SINEs and for SINEs harbouring an editing site in WT samples (Detected) or containing a differentially edited site. Results are shown as proportions of SINEs with the maximum count set to 100. See also Figure S6.

We further defined ‘differential’ sites as those that were ‘detected’ in the WT samples and showed higher (>2-fold) levels of editing than in *Adar1^mZα/mZα^* mice. 3,249 sites (8% of sites ‘detected’ in WT) were ‘differential’ and displayed ∼9% median editing in WT lungs, which was strongly reduced for the majority of sites in *Adar1^mZα/mZα^* mice (Figure 7B). Read depth was equivalent between ‘detected’ and ‘differential’ sites (data not shown), excluding the possibility that lack of editing in *Adar1^mZα/mZα^* samples was simply due to reduced sequence coverage. We then attempted to understand whether these Z*α* domain-dependent sites were characterised by unique properties. First, we annotated editing sites to 5’ untranslated regions (UTRs), coding sequences (CDSs), 3’UTRs, intronic and intergenic regions (Figure 7C). The majority of editing sites ‘detected’ in WT samples were found in intronic regions (∼83%). 3’UTRs accounted for ∼11% of edits and intergenic regions contained ∼5% of sites. Less than 1% of sites mapped to 5’UTRs and CDSs. The annotation of ‘differential’ sites was similar with a small increase in intronic sites (∼86%), while 8% of ‘differential’ edits were found in 3’UTRs.

Given that many A-to-I RNA editing events are known to occur in repetitive elements (REs) (Eisenberg and Levanon, 2018), we analysed the enrichment of editing events in WT samples, by computing observed versus expected numbers, for each RE class (Figure 7D). This analysis showed that editing events were greatly enriched within SINEs (Figure 7D) and further showed that, with the exception of the B4 family, all SINE sub-families exhibited enrichment, albeit to varying degrees (Figure S6A). Furthermore, there were no large-scale differences between the SINEs represented when comparing ‘detected’ and ‘differential’ sites (Figure S6B), suggesting that all SINEs had equal potential to contribute sites that were differentially edited between WT and *Adar1^mZα/mZα^* mice.

We next analysed the genomic distance of edited SINEs to another SINE in inverted orientation. In human, transcripts spanning inverted repeat *Alu* elements have been suggested to form duplex RNA structures recognised by ADAR1 and MDA5 (Ahmad et al., 2018; Mehdipour et al., 2020). Compared to all SINEs, we found that edited SINEs were closer to another SINE in inverted orientation (Figure 7E), although there was no difference between SINEs containing ‘detected’ (in WT samples) and ‘differential’ sites. Taken together, these data demonstrate that a subset consisting of about 8% of editing sites required a functional ADAR1-p150 Z*α* domain for efficient A-to-I conversion.

## Discussion

Although Z-nucleic acids were discovered ∼40 years ago, little is known to this date about their biological activities. This is in part due to their thermodynamic properties: the B- and A-conformations of dsDNA and dsRNA, respectively, are energetically favoured compared to the Z-conformation, making Z-DNA and Z-RNA difficult to study (Herbert, 2019). The formation of Z-DNA has been proposed to release torsional strain induced by the movement of polymerases (Wittig et al., 1992; Wolfl et al., 1995). Physiological functions of Z-RNA have remained enigmatic until recently. We and others proposed that Z-RNA is recognised by ZBP1 in settings of viral infection and autoinflammation, resulting in the induction of programmed cell death (Devos et al., 2020; Jiao et al., 2020; Maelfait et al., 2017; Sridharan et al., 2017; Wang et al., 2020; Zhang et al., 2020b).

Here, we studied ADAR1-p150 that like ZBP1 contains a Z*α* domain specialised in binding to Z nucleic acids. We report spontaneous induction of type I IFNs *in vivo* upon introduction of mutations into the ADAR1-p150 Z*α* domain that prevent binding to Z-DNA/RNA. This effect was observed in multiple organs and cell types from *Adar1^mZα/mZα^* mice and required MAVS. These data show that RIG-I-like receptors were activated by endogenously generated Z-RNAs, and that editing of these Z-RNAs by ADAR1-p150 limited the response. We therefore reveal type I IFN induction as a new biological function of Z-RNA.

Our computational analysis mapped ∼40,000 editing sites in lung RNA samples from WT mice. These sites were enriched in SINEs but not in other classes of REs. Furthermore, we found that edited SINEs were more likely to be in proximity to another SINE in inverted orientation. These results agree with previous findings demonstrating that editing sites are enriched in non-coding sequences containing self-complementary regions predicted to form duplex RNA structures (Eisenberg and Levanon, 2018; Porath et al., 2017; Solomon et al., 2017; Tan et al., 2017).

Importantly, we revealed that ∼8% of editing sites detected in WT samples required a functional ADAR1-p150 Z*α* domain. This subset of Z*α*-dependent sites mapped to genomic features and REs similarly to all detected sites. To characterise whether the sequences surrounding Z*α*-dependent editing sites have a propensity to form Z-RNA, we analysed sequences 500 nt up- and down-stream of editing sites. We tested GC content, given that GC-repeats have a higher tendency to adopt the Z-conformation (Davis et al., 1986). There was no enrichment of GC dinucleotides, or other motifs, around differentially edited sites compared to all detected sites (data not shown). We also used the Z-hunt algorithm (Champ et al., 2004; Ho et al., 1986) to predict the likelihood of sequences around editing sites to form the Z-conformation. In parallel, we calculated the distance of editing sites to genomic regions predicted by SIBZ to form the Z-conformation (Zhabinskaya and Benham, 2011). Neither computational approach revealed differences between ‘detected’ and ‘differential’ sites. It is noteworthy that both algorithms were developed for dsDNA and may be unsuitable for predicting the Z-conformation in RNA. Future studies will be required to dissect the properties the Z*α*-dependent RNA editing sites.

We also found diminished editing in WT samples at ∼14% of the sites detected as edited in *Adar1^mZα/mZα^* mice (data not shown). ADAR1-p150 is encoded by an ISG and *Adar1* transcript levels increased by 1.6-fold in *Adar1^mZα/mZα^* lungs (data not shown). Thus, it is likely that type I IFN-induced expression of ADAR1-p150 in mutant mice and recruitment of A-form dsRNAs via the dsRBDs explains editing at these sites.

*ADAR1* mutations in human cause AGS and include the missense p.Pro193Ala mutation in the Z*α* domain. Interestingly, this *ADAR1* allele is common with frequencies of up to ∼1/160 (www.ensembl.org) (Mannion et al., 2014). In AGS patients, homozygous p.Pro193Ala mutation has not been observed; instead, this mutation occurs together with other *ADAR1* mutations (Rice et al., 2012). It is therefore likely that *ADAR1* p.Pro193Ala is hypomorphic and does not cause disease when present homozygously. Consistent with this notion, we found that *Adar1^mZα/mZα^* mice did not have any gross abnormalities and were fertile. These mice nonetheless displayed type I IFN and ISG induction in multiple organs. This included the lung and bestowed protection against IAV infection at early stages. It is therefore conceivable that the *ADAR1* p.Pro193Ala variant has been maintained in humans by providing a selective advantage during viral infections due to elevated expression of antiviral factors at baseline.

It is noteworthy in this context that anti- and pro-viral functions of ADAR1 have been reported. ADAR1 limits or controls replication of several viruses, including measles virus, members of *Paramyxoviridae* family, IAV, HIV-1, vesicular stomatitis virus and Hepatitis delta virus (Casey, 2012; Li et al., 2010; Vogel et al., 2020; Ward et al., 2011; Weiden et al., 2014). However, for other viruses such as Zika and KSHV, ADAR1 has been shown to facilitate replication (Zhang et al., 2020a; Zhou et al., 2019). It will be interesting for future studies to determine the role of the *ADAR1* variants unable to bind Z nucleic acids in these viral infections.

ADAR1 has recently emerged as a promising target for cancer treatment. If expressed by transformed cells *in vitro* or by tumours *in vivo*, ADAR1 protects against both cell death and anti-cancer immune responses (Gannon et al., 2018; Ishizuka et al., 2019; Liu et al., 2019). Loss of ADAR1 in cancer cells results in death or reduced growth and sensitises to immunotherapy. Interestingly, the protective effects appear to depend on ADAR1-p150 (Gannon et al., 2018; Ishizuka et al., 2019). It is therefore possible that endogenous Z-RNAs induce anti-cancer effects upon ADAR1 loss. Future studies should test this, for example by reconstitution of ADAR1-p150 mutants unable to bind Z-RNA. Furthermore, development of inhibitors that target the Z*α* domain of ADAR1, the Z*α* - Z-RNA interaction or Z-RNA formation should be considered. Compared to deaminase inhibitors, such ‘Z-inhibitors’ would have the advantage of specifically targeting the p150 isoform, avoiding possible detrimental consequences of targeting ADAR1-p110 (Pestal et al., 2015).

In addition to the activation of MDA5, loss of ADAR1 also results in activation of the OAS-RNaseL system and PKR. The latter may be particularly important in cancer settings (Gannon et al., 2018; Ishizuka et al., 2019; Liu et al., 2019). Although PKR activation results in a global shutdown of translation, some proteins are selectively made and mediate the integrated stress response (Pakos-Zebrucka et al., 2016). These include the transcription factor ATF4 that induces stress-response genes. We observed moderate induction of some ATF4-dependent genes (Harding et al., 2003) including *Asns* (1.3-fold), *Slc7a5* (1.3-fold), *Slc7a11* (1.9-fold) and *Mthfd2* (1.7-fold) (data not shown). It is therefore possible that editing of Z-RNA by ADAR1-p150 limits not only type I IFN induction but also PKR-dependent stress responses.

In conclusion, we discovered MAVS-dependent type I IFN induction as a biological function of Z-RNA that is curtailed by ADAR1-p150. These insights are not only of fundamental value but also have important implications for understanding and modulating detrimental and beneficial type I IFN responses in autoinflammation and cancer.

## Materials and Methods

### Mice

*Adar1^+/fl-mZα^* mice were generated by Cyagen. In brief, genomic fragments containing homology arms were amplified from a BAC and were sequentially assembled into a targeting vector together with recombination sites and selection markers as shown in Figure S1A. Successful assembly of the targeting vector was verified by restriction digest and sequencing. The linearised vector was subsequently delivered to ES cells (C57BL/6) via electroporation, followed by drug selection, PCR screening and sequencing. After confirming correctly targeted ES clones via Southern blotting, we selected clones for blastocyst microinjection, followed by chimera production. Founders were confirmed as germline-transmitted via crossbreeding with WT animals. The Neo cassette was flanked by Rox sites and contained a Dre recombinase controlled by a promoter active in the germline, resulting in deletion of the Neo cassette in F1 animals (Figure S1A). These *Adar1^+/fl-mZα^* mice were further crossed with *Pgk-Cre* mice provided by Samira Lakhal-Littleton to produce *Adar1^+/mZα^* animals (Figure S1A). The following PCR primers were used for genotyping:

F, 5’-TGACGAGAGACTTGTTTTCCTAGCATG-3’,

R1, 5’-TGCCTCAATGAGACCTCCAACTTAACTC-3’,

R2^WT^, 5’-CAGGGAGTACAAAATACGATT-3’, and

R2^MUT^, 5’-CAGGGAGGCCAAAATACGAGC-3’.

PCR with primers F and R1 yielded a product of 357 bp for the WT *Adar1* allele and a 421 bp product for both ‘*fl-mZα*’ and ‘*mZα*’ alleles. PCR with primers F and R2^WT^ resulted in 1095 and 1158 bp products for the WT and ‘fl*-mZα*’ alleles, respectively, and no product for the ‘*mZα*’ allele. Finally, PCR with primers F and R2^MUT^ resulted in a 1158 bp product for the ‘*mZα*’ allele only.

*Μavs^-/-^* mice were a gift from C. Reis e Sousa and were originally from J. Tschopp (Michallet et al., 2008). All mice were on the C57BL/6 background. This work was performed in accordance with the UK Animal (Scientific Procedures) Act 1986 and institutional guidelines for animal care. This work was approved by project licenses granted by the UK Home Office (PPL numbers PC041D0AB, PBA43A2E4 and P79A4C5BA) and was also approved by the Institutional Animal Ethics Committee Review Board at the University of Oxford.

### Cell culture

Lung fibroblasts and MEFs were grown in DMEM and BMMCs in RPMI, as described previously (Li et al., 2013; Maelfait et al., 2017). Media were supplemented with 10% heat-inactivated FCS and 2 mM L-glutamine; for BMMCs, 200 U/ml recombinant mouse GM-CSF (Peprotech) was added. MEFs were cultured at 3% oxygen.

### RNA extraction and RT-qPCR

Organs collected from freshly killed mice (8-10 weeks of age) were snap frozen in liquid nitrogen immediately after dissection and stored at −80°C until further processing. Organs were homogenised with glass beads (425-600 μm, Sigma-Aldrich) in TRIzol (Thermo Fisher Scientific) using a FastPrep F120 instrument (Thermo Savant). RNA was extracted following the manufacturer’s instructions and further purified using RNeasy Plus columns (Qiagen) including a gDNA eliminator column step. cDNA synthesis was performed with SuperScript II reverse transcriptase (Thermo Fisher Scientific) with random hexamer (Qiagen) or oligo (dT)12-18 (Thermo Fisher Scientific) as primers. qPCR was done using Taqman Universal PCR Mix (Thermo Fisher Scientific) and Taqman probes (Applied Biosystems). Alternatively, qPCR was performed using EXPRSS SYBR GreenER qPCR Supermix (Thermo Fisher Scientific) and DNA oligonucleotides (Sigma Aldrich). qPCR was performed on a QuantStudio 7 Flex real-time PCR system (Applied Biosystem). The qPCR probes and primers used in this study are listed in Table S2.

### *In vivo* infection

WT and *Adar1^mZα/mZα^* mice were used at 8-10 weeks of age. Mice were intranasally inoculated with 50 μl X31 (0.04 haemagglutination units (HAU)) diluted in viral growth medium (VGM; DMEM with 1% bovine serum albumin (Sigma-Aldrich A0336), 10 mM HEPES buffer, penicillin (100 U/ml) and streptomycin (100 μg/ml)) or mock infected with 50 μl VGM under light isoflurane anaesthesia. Animals were assessed daily for weight loss and signs of disease. Mice reaching 20% weight loss were euthanised.

### Western Blot

Cells were lysed in RIPA buffer (50 mM Tris.HCl, pH7.4; 150 mM NaCl; 1% NP-40 (Sigma-Aldrich), 0.5% Deoxycholate, 0.1% SDS and Complete protease inhibitor (Roche)) at 4°C for 10 minutes. Protein lysates were then cleared by centrifugation at 13000 rpm for 10 minutes. Samples were mixed with NuPAGE SDS-PAGE sample loading buffer (ThermoFisher) containing 10% 2-mercaptoethanol. A primary antibody against ADAR1 was purchased from Santa Cruz (sc-73408). The antibody recognising ISG15 was a gift from Klaus-Peter Knobeloch. HRP-coupled secondary antibodies were from GE Healthcare.

### Flow cytometry

Lungs from 8-10 week old mice were dissected and mechanically disrupted using scissors before incubation in RPMI containing 1 μg/ml type II collagenase (Worthington Biochemical Corporation) and 40 U/ml DNase I (Sigma Aldrich) at 37°C for 60 minutes, with resuspension after 30 minutes to facilitate tissue dissociation. Cells were filtered through a 70 μm cell strainer (BD Falcon), rinsed with RPMI and pelleted at 400 x g for 5 minutes. The cell pellet was resuspended in 5 ml RBC lysis buffer (Qiagen), incubated at room temperature for 5 minutes and then washed twice with 45 ml RPMI. Cells were resuspended in 500 μl FACS buffer (PBS containing 10% (v/v) FCS and 2 mM EDTA) and passed through a 70 μm cell strainer. Viable cells were counted using a haemocytometer. Cells were washed with PBS before incubation with LIVE/DEAD Fixable Aqua Dead Cell Stain (Invitrogen) diluted 1:200 in PBS for 30 minutes at room temperature. Cells were washed once with FACS buffer and then stained with surface antibodies diluted 1:200 in Brilliant buffer (BD Biosciences) for 30 minutes. Cells were sorted directly into TRIzol-LS Reagent (Thermo Fisher Scientific) on BD FACSAria II and III machines (BD Biosciences). Alternatively, 1.5×10^6^ cells were stained and analysed using an Attune NxT Flow Cytometer (Thermo Fisher Scientific). Data were analysed using FlowJo (v10.6.2).

### Magnetic cell fraction

Cells from lungs were prepared as described above. 10^7^ cells were resuspended in 90 μl of MACS buffer (PBS containing 0.5% BSA and 2 mM EDTA) and then incubated with 10 μl of CD45 microbeads (Miltenyi) for 15 minutes. The mixture was then washed with MACS buffer and resuspend in 500 μl MACS buffer for magnetic separation on MACS LS columns (Miltenyi) according to the manufacturer’s instructions. Cells were recovered from the flow-through (CD45-) and column (CD45+). 10% of cells were stained and analysed by FACS to confirm purity. The remaining cells were pelleted, resuspend in TRIzol (ThermoFisher) and processed for RT-qPCR.

### Bone marrow chimera

B6.SJL-CD45.1 mice were used as bone marrow recipients and were lethally irradiated twice (4.5 Gy for 300 seconds, separated by a ∼3 hour rest). Mice were then injected intravenously with bone marrow from either WT (CD45.2) or *Adar1^mZα/mZα^* mice. Recipient mice received antibiotics (0.16 mg/mL, Enrofloxacin (Baytril), Bayer Corporation) in drinking water for four weeks following irradiation and were rested for >8 weeks before tissue collection.

### RNA-seq and data processing

Stranded Illumina sequencing libraries were prepared with the RNA-Seq Ribozero kit from isolated RNAs and submitted for PE150 sequencing using an Illumina NovaSeq6000 machine, yielding ∼100M reads per sample. Sequencing data was processed using a Nextflow v20.07 (Di Tommaso et al., 2017) pipeline automating quality control using FastQC v0.11.8 (bioinformatics.babraham.ac.uk/projects/fastqc/), quality and adapter trimming using cutadapt v1.18 (Martin, 2011), contaminant detection using screen.sh (within BBMap v36.20, sourceforge.net/projects/bbmap/), strand-aware alignment using HISAT2 v2.1.0 (Kim et al., 2019) and STAR v2.7.1a (Dobin et al., 2013), post-alignment quality-assurance using ‘gene body coverage’, ‘transcript integrity’, and ‘inner distance’ metrics from RSeQC v2.6.4 (Wang et al., 2012), and strand-specific counting of uniquely-mapping reads using featureCounts (within Subread v1.6.4, (Liao et al., 2014)) against Ensembl GRCm38.100 annotations. Additional, unstranded counts were obtained with featureCounts against a database of repetitive elements previously prepared for GRCm38 (Attig et al., 2017) using reads unassigned to features during the previous step.

### Differential expression analysis

Downstream differential expression analysis was conducted using counts obtained for STAR read mappings using DESeq2 (Love et al., 2014) (v1.22.1) within R (v4.0.2). Gene ontology analysis was performed using goseq (Young et al., 2010) (1.34.1). Heatmaps were generated using the pheatmap package (v1.0.12).

### Detection of A-to-I editing

A Python 3.8 program, edIted (github.com/A-N-Other/pedestal), was produced to identify editing sites. edIted performs stranded assessments of RNA editing from samtools mpileup (Li, 2011) data, building on the Dirichlet-based models implemented in ACCUSA2 and JACUSA (Piechotta and Dieterich, 2013; Piechotta et al., 2017). When run with test data alone, edIted runs in ‘detect’ mode, finding base modifications by comparing the goodness of fit of Dirchlet models of the base error (derived from the Phred quality data in the mpileup input) and the background sequencing error to the base frequencies recorded at a specific position. With an additional control dataset, edIted runs in ‘differential’ mode, performing the above analysis to determine significantly edited sites before additionally testing for differential editing by comparing the goodness of fit of Dirichlet models of the base error from the test and control datasets to their own and each other’s base frequencies. When biological replicates are provided, edIted adjusts the reported Z scores to reflect the proportion of test dataset samples displaying editing. edIted was run in both modes with samtools mpileup files (supplemented with TS tag metadata) separately for HISAT2 and STAR alignments of the data with the flags ‘-- min_depth 5 --min_alt_depth 2 --min_edited 0.01 --max_edited 0.9 --z_score 2.58’. For differential analyses the ‘--min_fold 2’ flag was used and, where considering biological replicates the ‘--reps 3’ flag, such that editing is required in all three samples. All analyses were conducted supplying BED files of ENCODE blacklisted regions (Amemiya et al., 2019) and known splice sites (regions set to splice site +/− 2 nts) to the ‘--blacklist’ flag. Sites that were found in common between the HISAT2- and STAR-mapped data were retained for further analysis.

### Analysis of A-to-I editing sites

Sites obtained from edIted were assigned to genomic features using annotatr v1.16 within R (Cavalcante and Sartor, 2017). Assessments of editing enrichment within repetitive elements were conducted using regioneR v1.22 within R (Gel et al., 2016) using randomization-based permutation tests with 100 bootstraps. Assessment of distance to neighbouring inverted SINE super-family (B1, B2, B3, B4) members was conducted with bedtools v2.29.2 closest (Quinlan and Hall, 2010) using the ‘-io -S’ flags. Outputs and the detailed statistics were produced with GraphPad Prism v8.

### ELISA

Mouse IL-6 was quantified by uncoated ELISA Kit (ThermoFisher) according to manufacturer’s instruction.

## Author contributions (using the CRediT taxonomy)

Conceptualisation: Q.T. and J.R.; Methodology: Q.T., R.E.R. and G.Y.; Software: R.E.R. and G.Y.; Validation: Q.T. and J.R.; Formal analysis: Q.T., R.E.R., G.Y. and J.R.; Investigation: Q.T., R.E.R., G.Y., A.K.H., T.K.T. and A.B.; Resources: A.T. and G.K.; Data curation: Q.T., R.E.R. and G.Y.; Writing – Original Draft: Q.T., G.Y. and J.R.; Writing – Review & Editing: all authors; Visualisation: Q.T., R.E.R., G.Y. and J.R.; Supervision: J.R., A.T., and G.K.; Project administration: Q.T.; Funding acquisition: J.R.

## Acknowledgments

The authors thank P. Shing Ho and Craig Benham for providing access to the Z-hunt and SIBZ codes, respectively. We further thank Daniel Stetson, Andrew Oberst, Jonathan Maelfait, Caetano Reis e Sousa, David Ron, Annemarthe van Der Veen and members of the Rehwinkel lab for discussion. The authors thank Ziqi Long and Oliver Bannard for their help with generating BM chimeras. This work was funded by the UK Medical Research Council [MRC core funding of the MRC Human Immunology Unit; J.R.], the Wellcome Trust [grant number 100954; J.R.], the Lister Institute [J.R.] and the Francis Crick Institute [G.K.], which receives its core funding from Cancer Research UK, the UK Medical Research Council, and the Wellcome Trust. T.K.T. was funded by the Townsend-Jeantet Charitable Trust (charity number 1011770) and the EPA Cephalosporin Early Career Researcher Fund. The funders had no role in study design, data collection and analysis, decision to publish, or preparation of the manuscript.

## Figures and figure legends

**Figure S1, related to Figure1.**
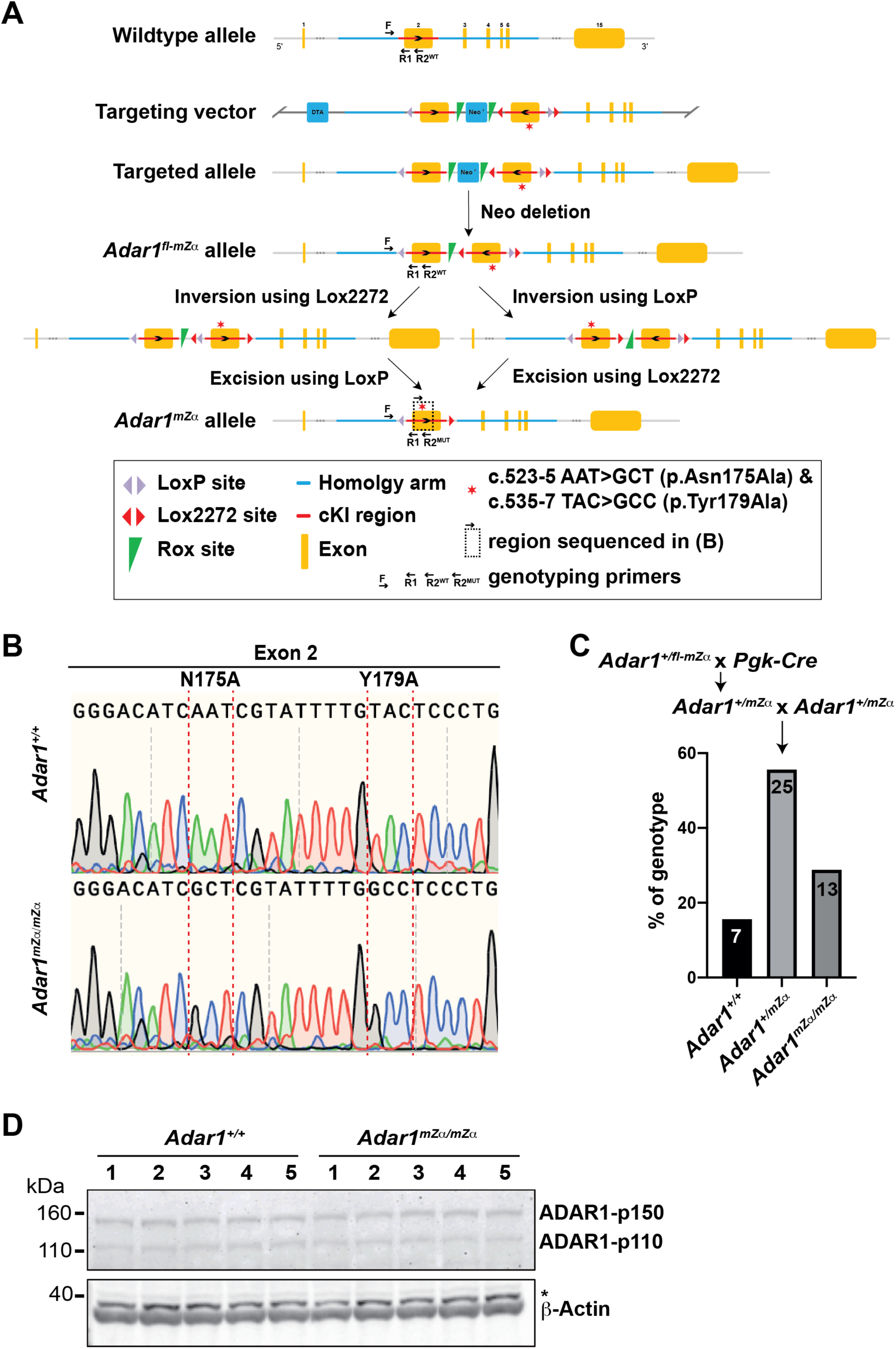
Generation of *Adar1^mZα/mZα^* animals. **A.** Schematic representation of the *Adar1* WT allele, targeting vector, targeted allele, *Adar1^fl-mZα^* allele and the two-step Cre-mediated recombination process that resulted in the *Adar1^mZα^* allele. Please see text for details. cKI, conditional knock-in. **B.** Genomic DNA was prepared from WT and *Adar1^mZα/mZα^* animals and the mutated region in exon 2 was sequenced. **C.** *Adar1^+/fl-mZα^* mice were bred with the *Pgk-Cre* line. *Adar^+/mZα^* offspring were then mated to generate *Adar1^mZα/mZα^* animals. The numbers and percentages of animals obtained with the indicated genotypes are shown. **D.** BMMCs were grown from bone marrow from five mice of the indicated genotypes. Protein extracts were used for western blot with *α*-ADAR1 antibody. *β*-Actin served as a loading control. *, non-specific band

**Figure S2, related to Figure 3.**
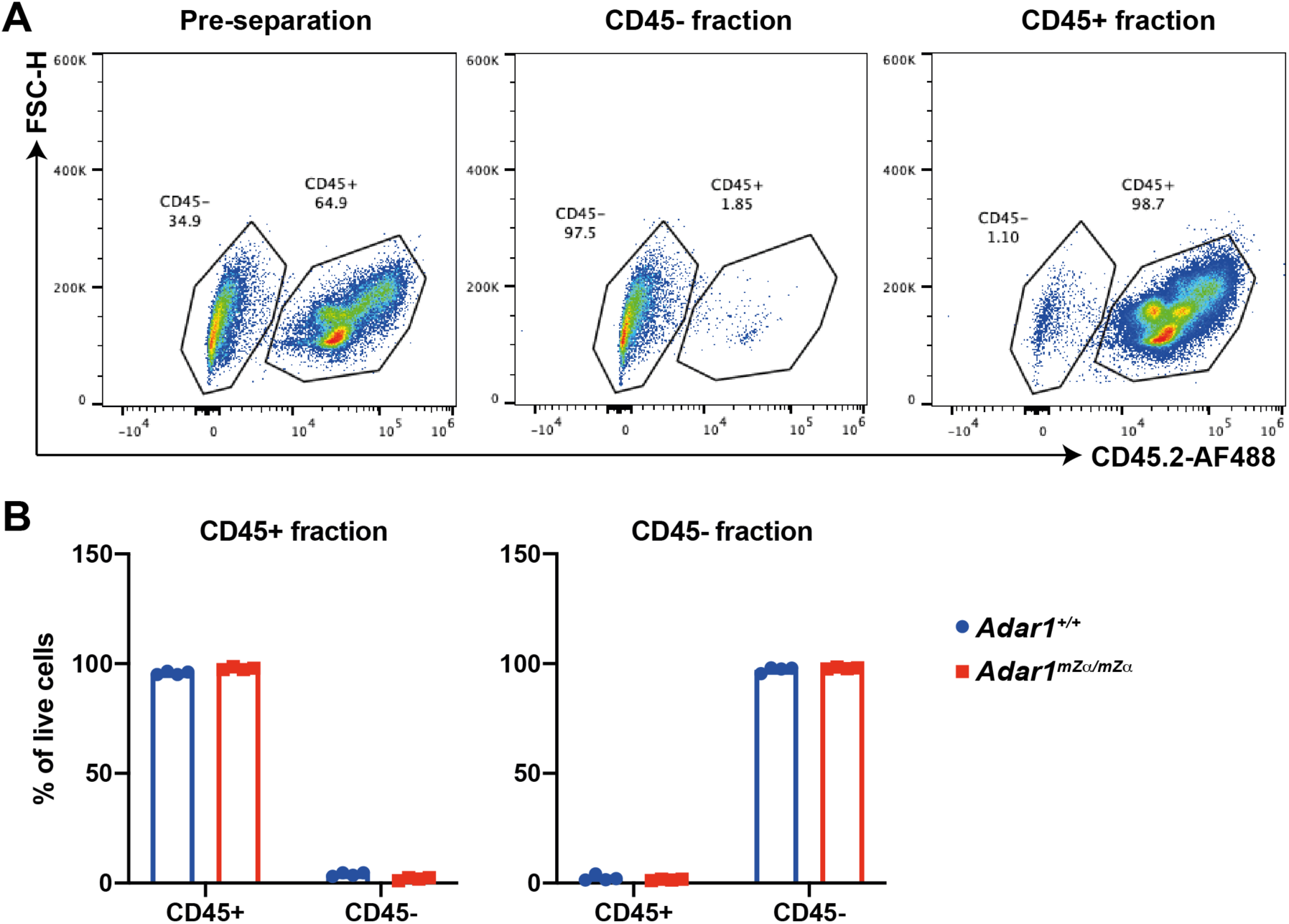
MACS separation of lung cells. **A.** Cell surface levels of CD45 were analysed by flow cytometry in single cell suspensions obtained from lung tissue before MACS (left) and in CD45- and CD45+ cell fractions obtained after MACS (middle and right). Data are from a representative WT animal. **B.** The percentage of CD45-expressing cells is shown for CD45+ and CD45-MACS fractions. Data points represent individual animals (n=4) from a representative experiment and bars indicate the mean.

**Figure S3, related to Figure 4A and 4B.**
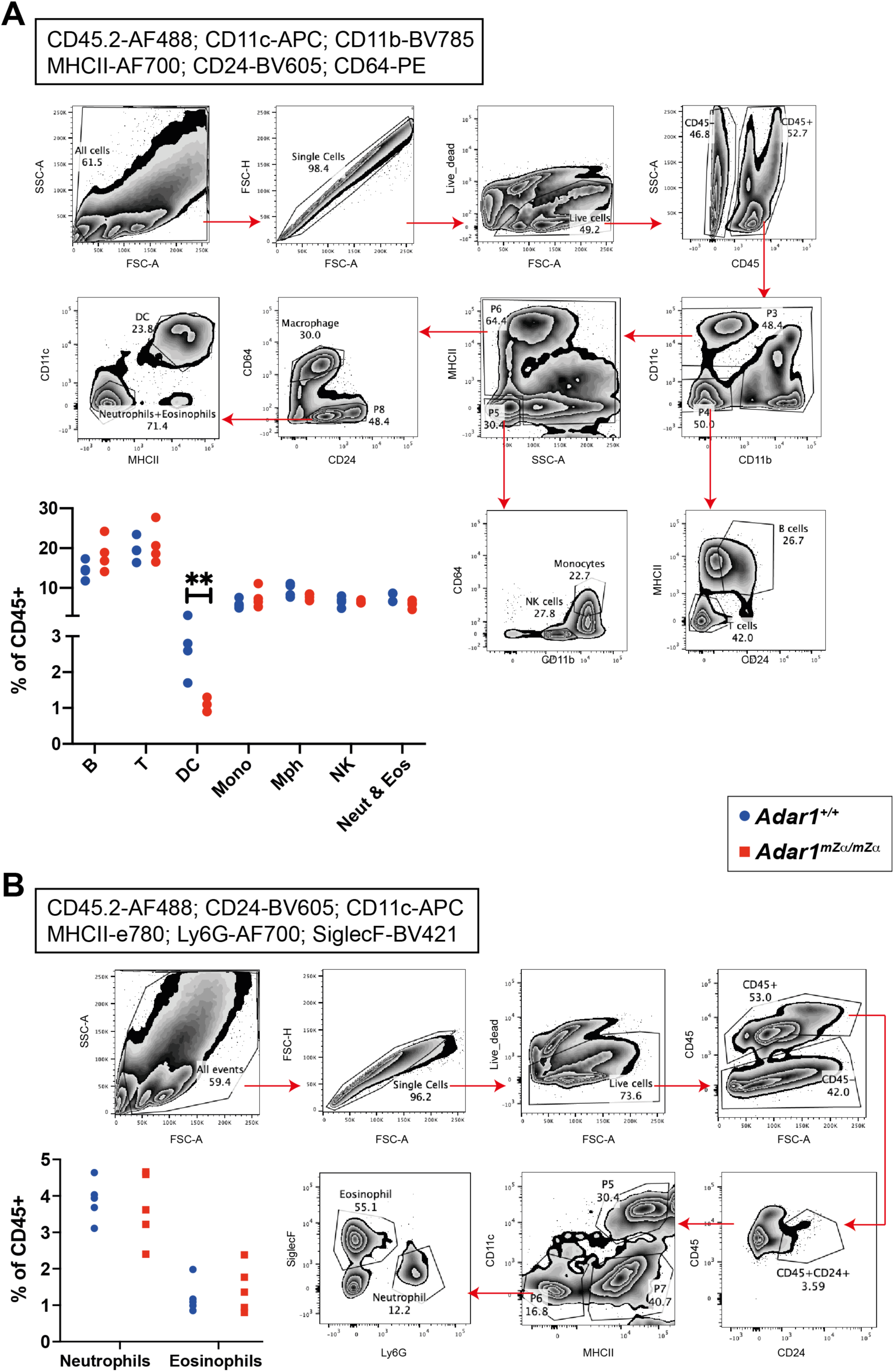
Gating strategy for sorting of haematopoietic lung cells. Two staining panels were used to identify and isolate haematopoietic cell populations by FACS. Panel (A) is related to Figure 4A and panel (B) to Figure 4B. Antibodies and conjugated fluorophores are shown in boxes. Gating strategies are shown for a representative WT (A) and *Adar1^mZα/mZα^* (B) animal. Bar graphs show the proportion of each cell population as a percentage of CD45+ cells. Each dot represents an individual mouse and data from two independent experiments were pooled (**p<0.01, unpaired t test).

**Figure S4, related to Figure 4C.**
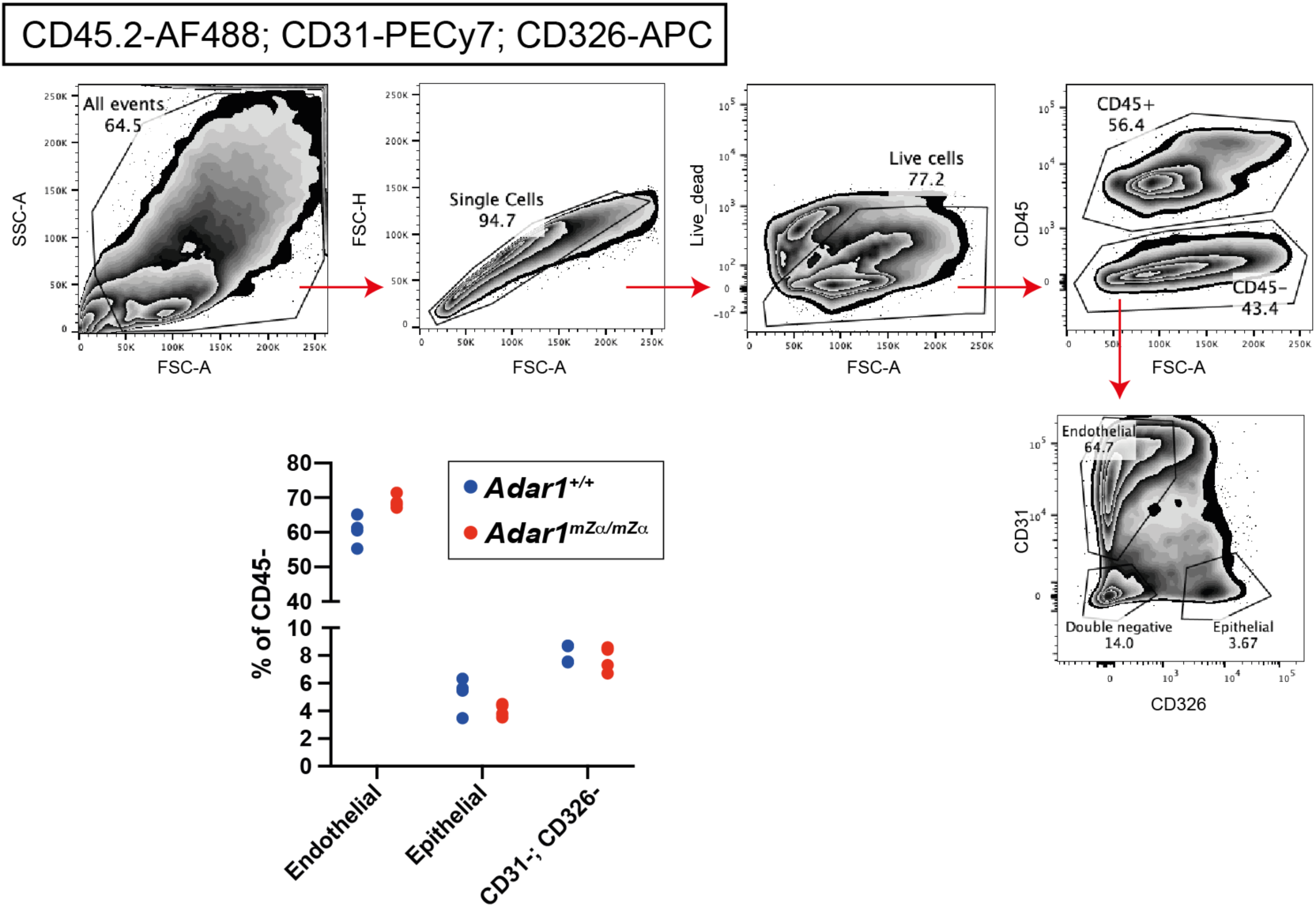
Gating strategy for sorting of stromal lung cells. The staining panel used to identify and isolate non-haematopoietic cell populations by FACS is shown. Antibodies and conjugated fluorophores are shown in the box. The gating strategy is shown for a representative WT animal. The bar graph shows the proportion of each cell population as a percentage of CD45-cells. Each dot represents an individual mouse.

**Figure S5, related to Figure 4D.**
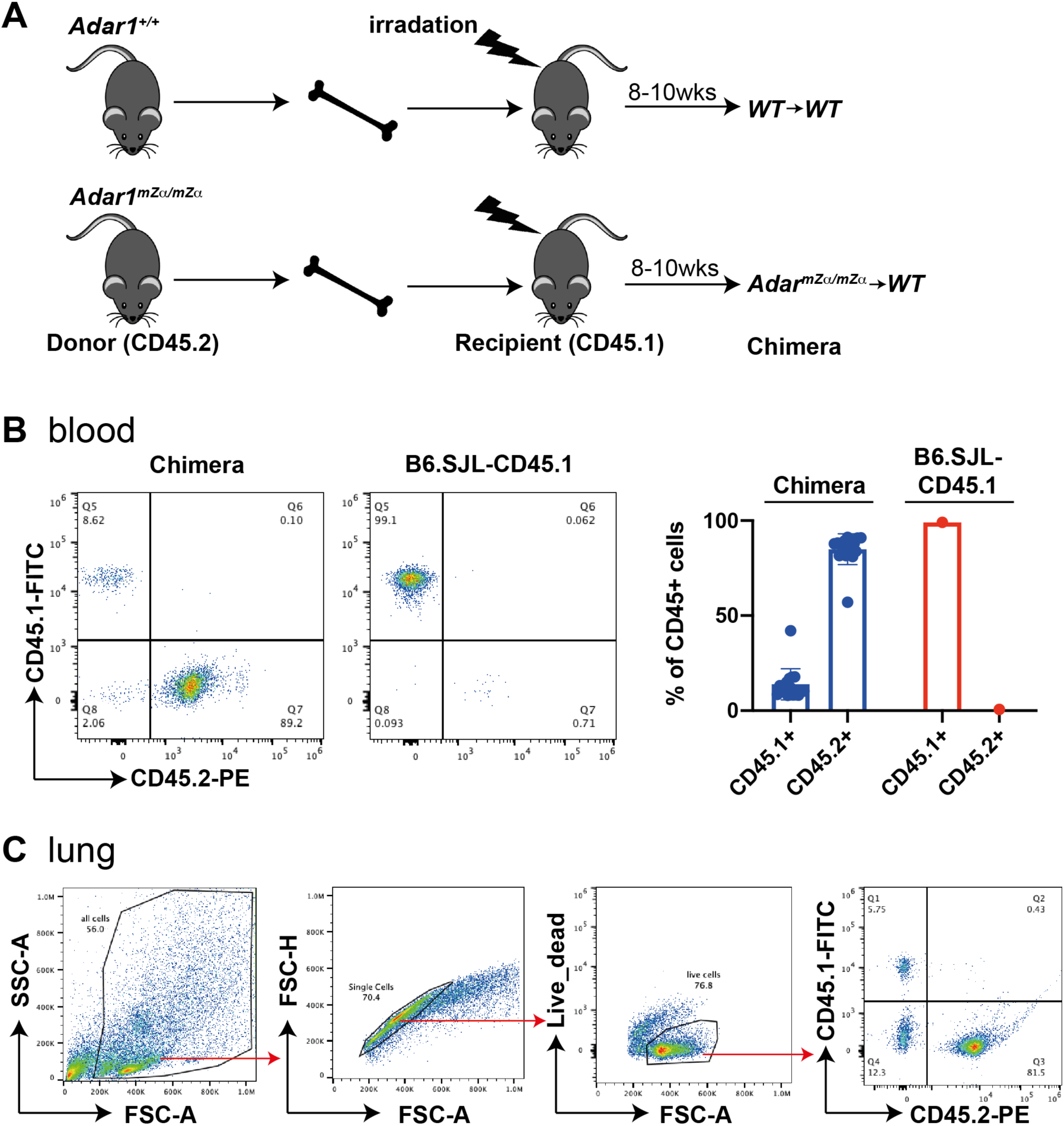
Analysis of BM chimeras. **A.** Schematic representation of the generation of BM chimeric animals. **B.** White blood cells from BM chimeric mice and, as control, from an untreated B6.SJL-CD45.1 animal, were analysed by FACS. Cells were gated on single, live cells. Representative FACS plots from a WT→WT animal (left) and pooled data from two independent experiments involving a total seven WT→WT and eight *Adar1^mZα/mZα^*→WT BM chimeric animals (right) are shown. Bars show the mean and error bars represent SD. **C.** Lung cells from BM chimeric mice were analysed by FACS. Data from a representative WT→WT animal are shown.

**Figure S6.**
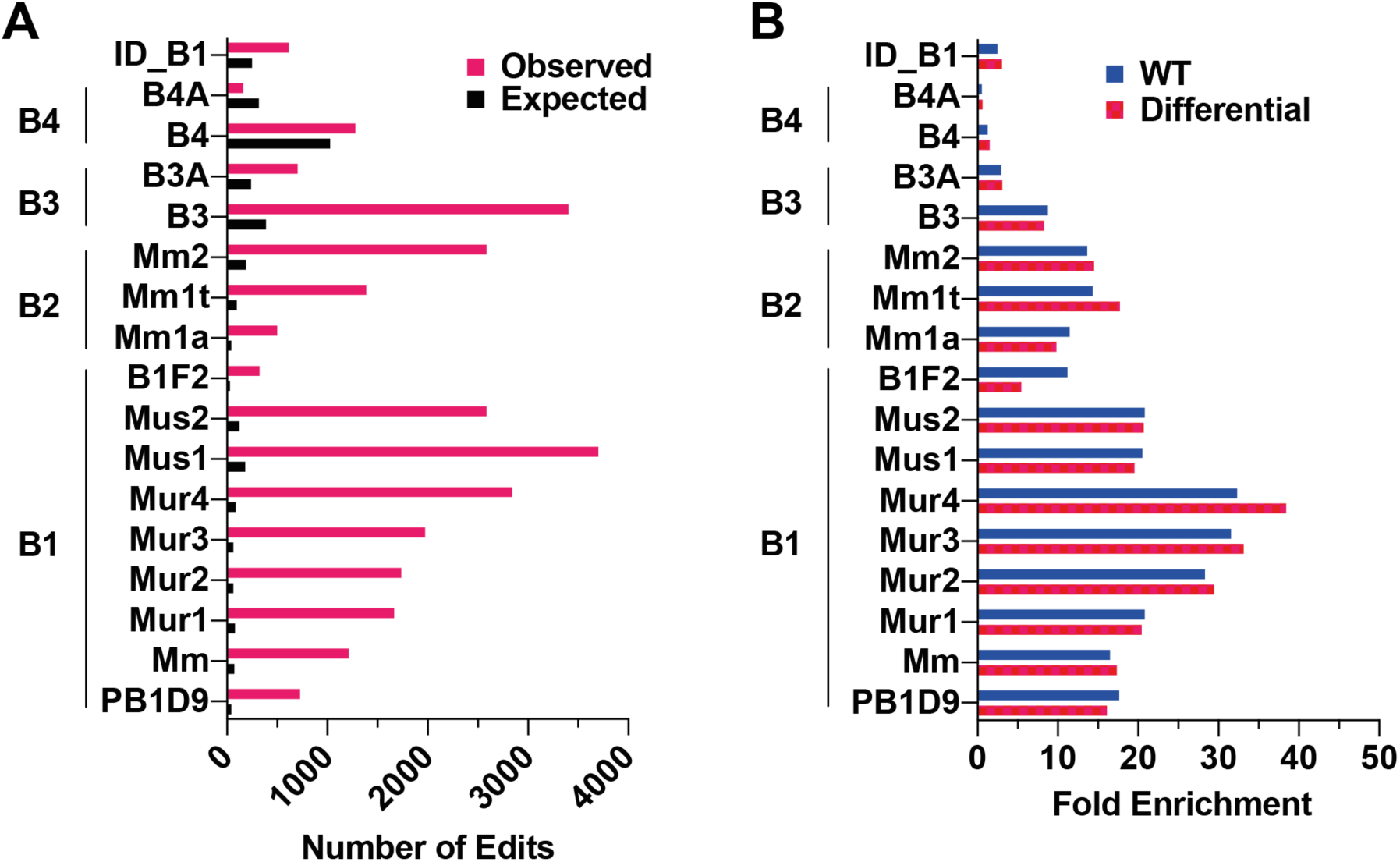
Analysis of RNA editing in SINEs. Related to Figure 7. **A.** Sub-families of SINEs were analysed as in Figure 6D, but are plotted to include those where observed or expected values exceeded 250. **B.** Fold enrichments of editing events were calculated relative to expected numbers of edits for SINE sub-families using editing events detected in WT samples and differentially edited sites.

**Table S1.**
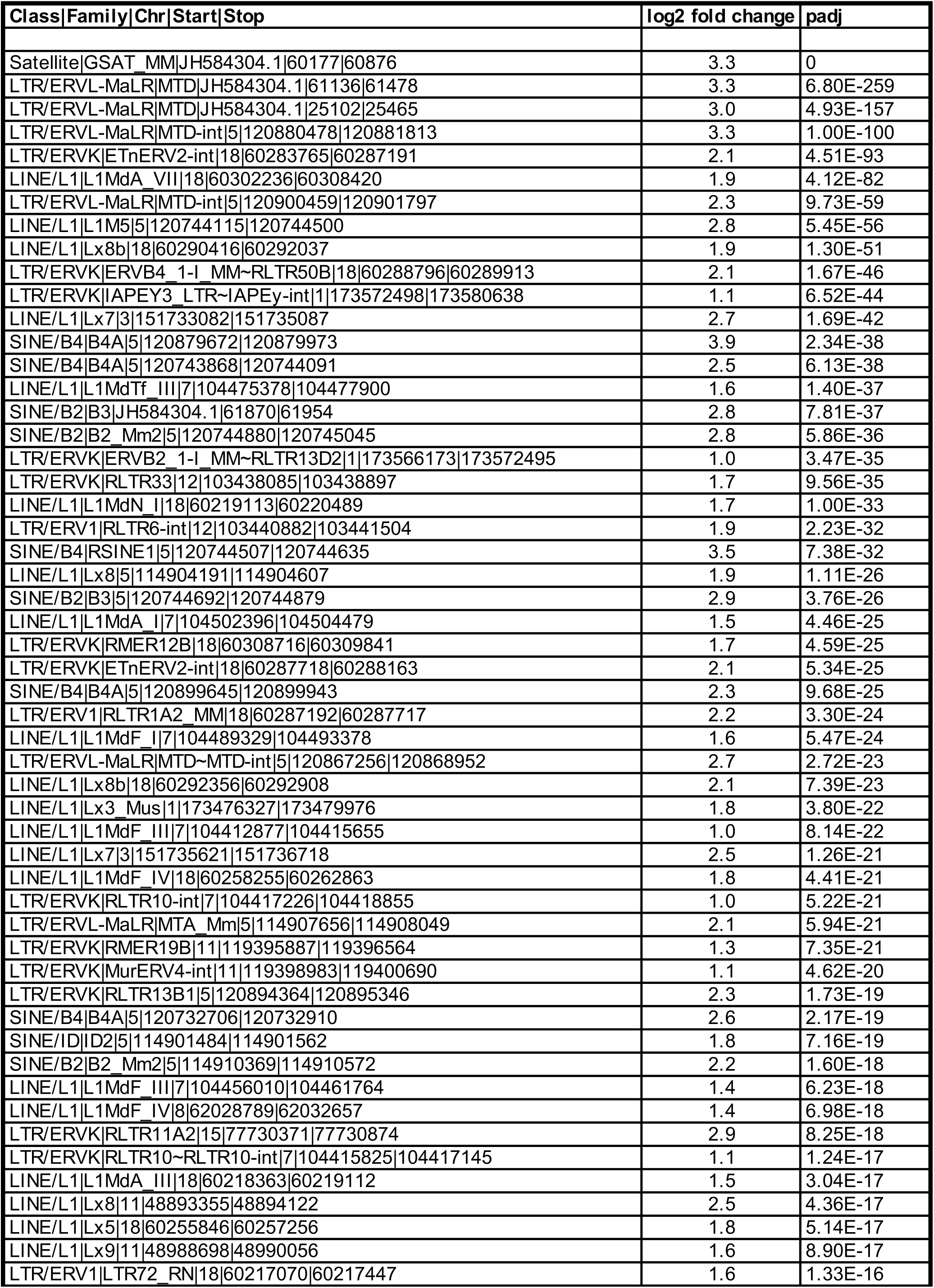
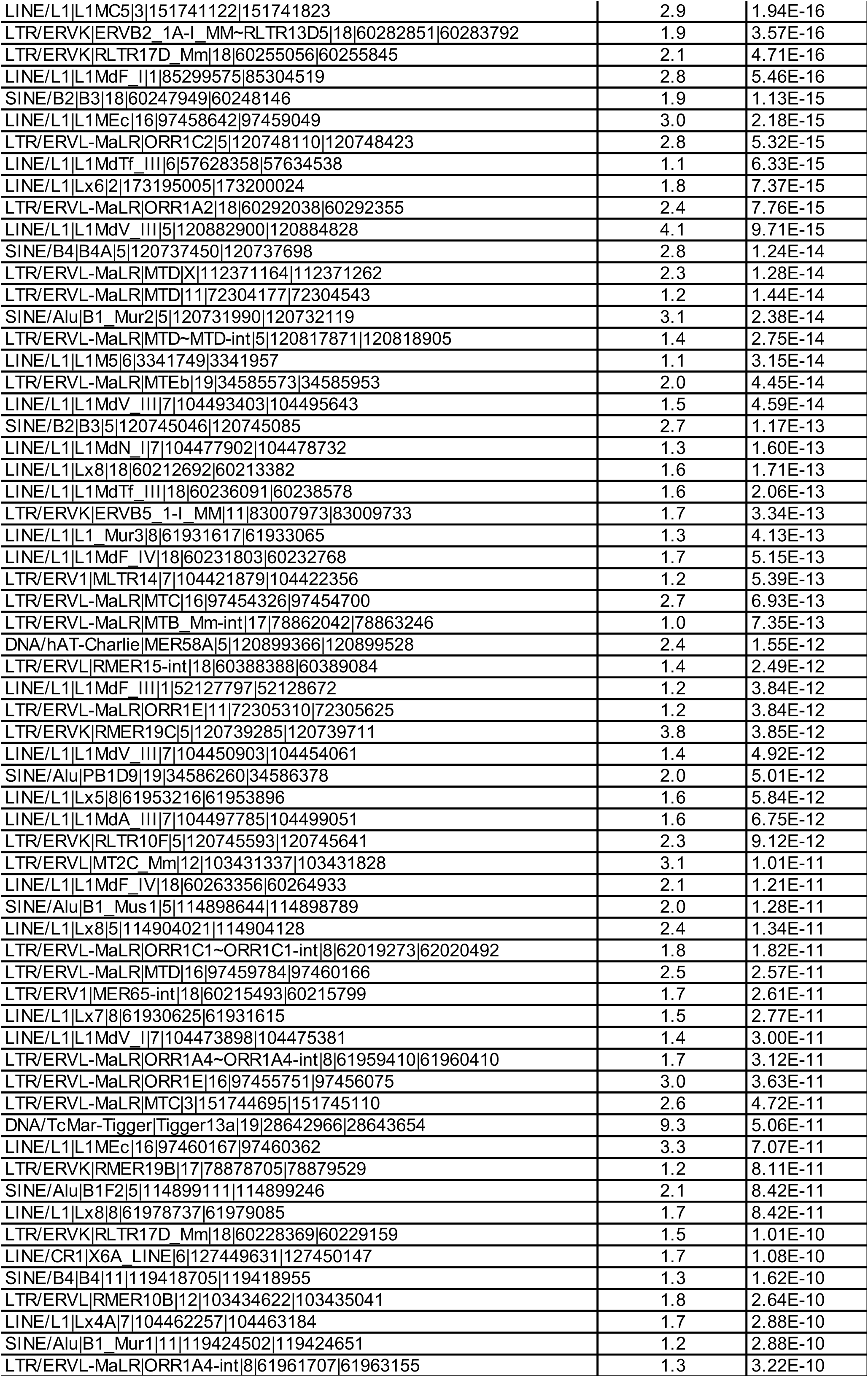
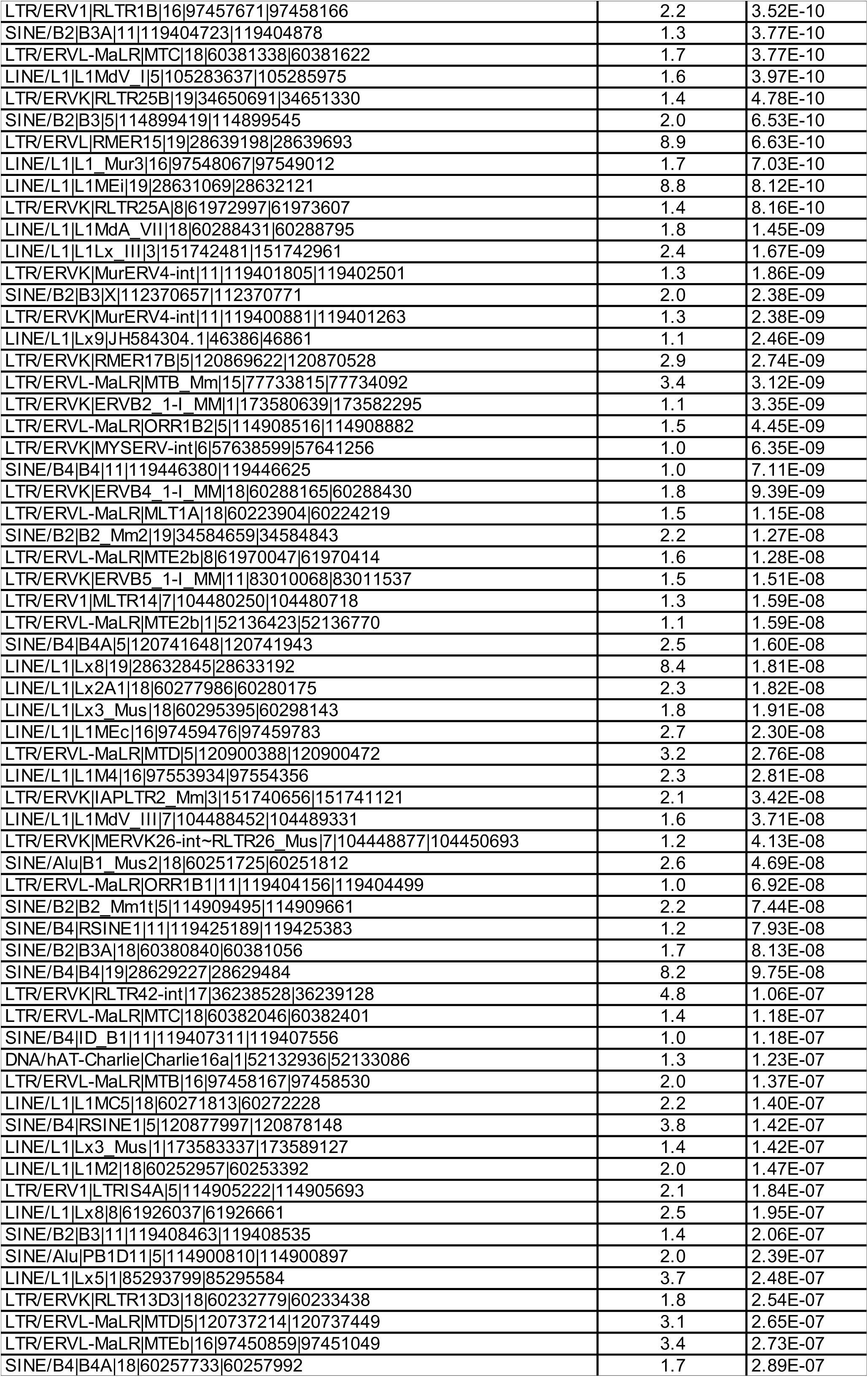
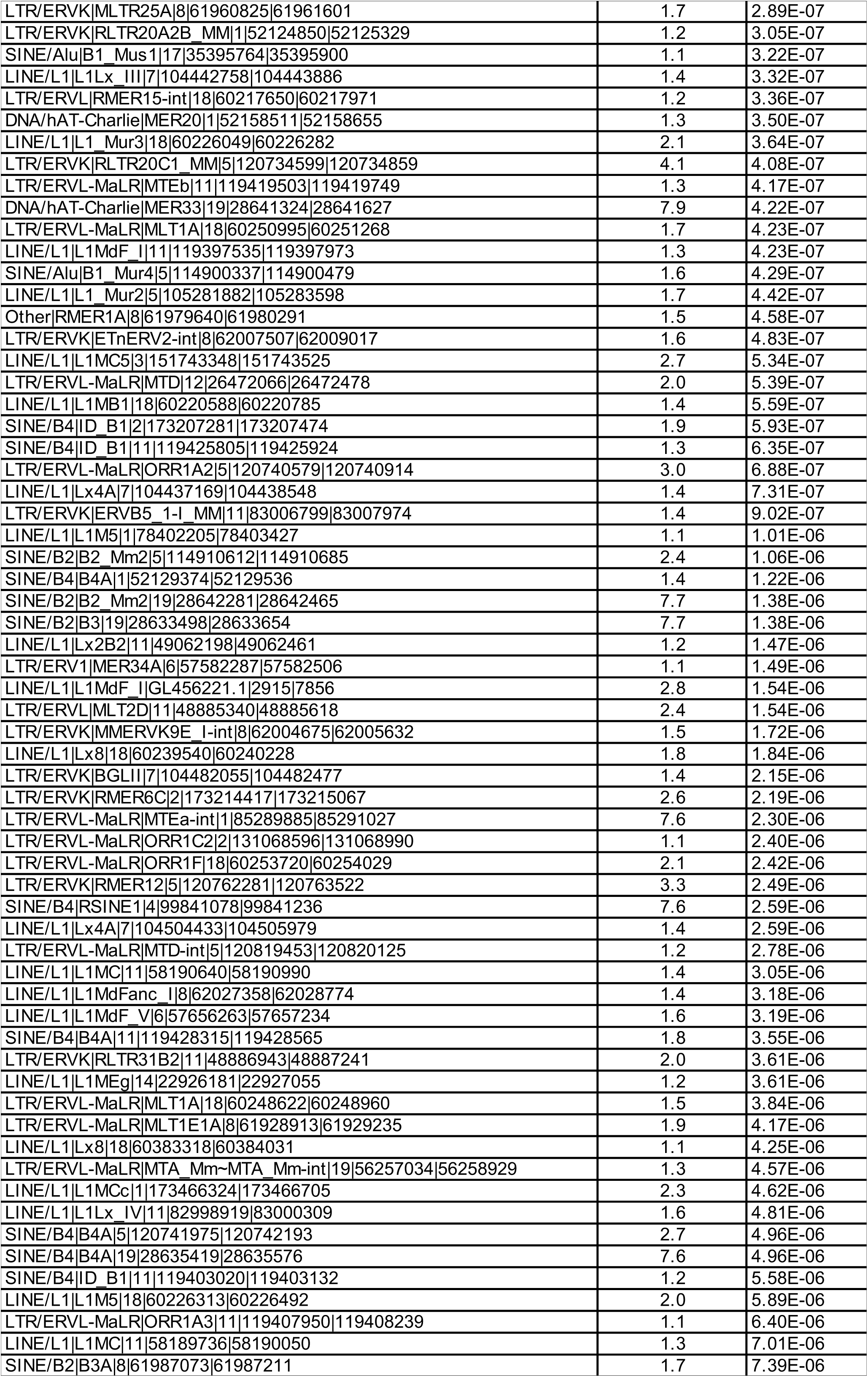
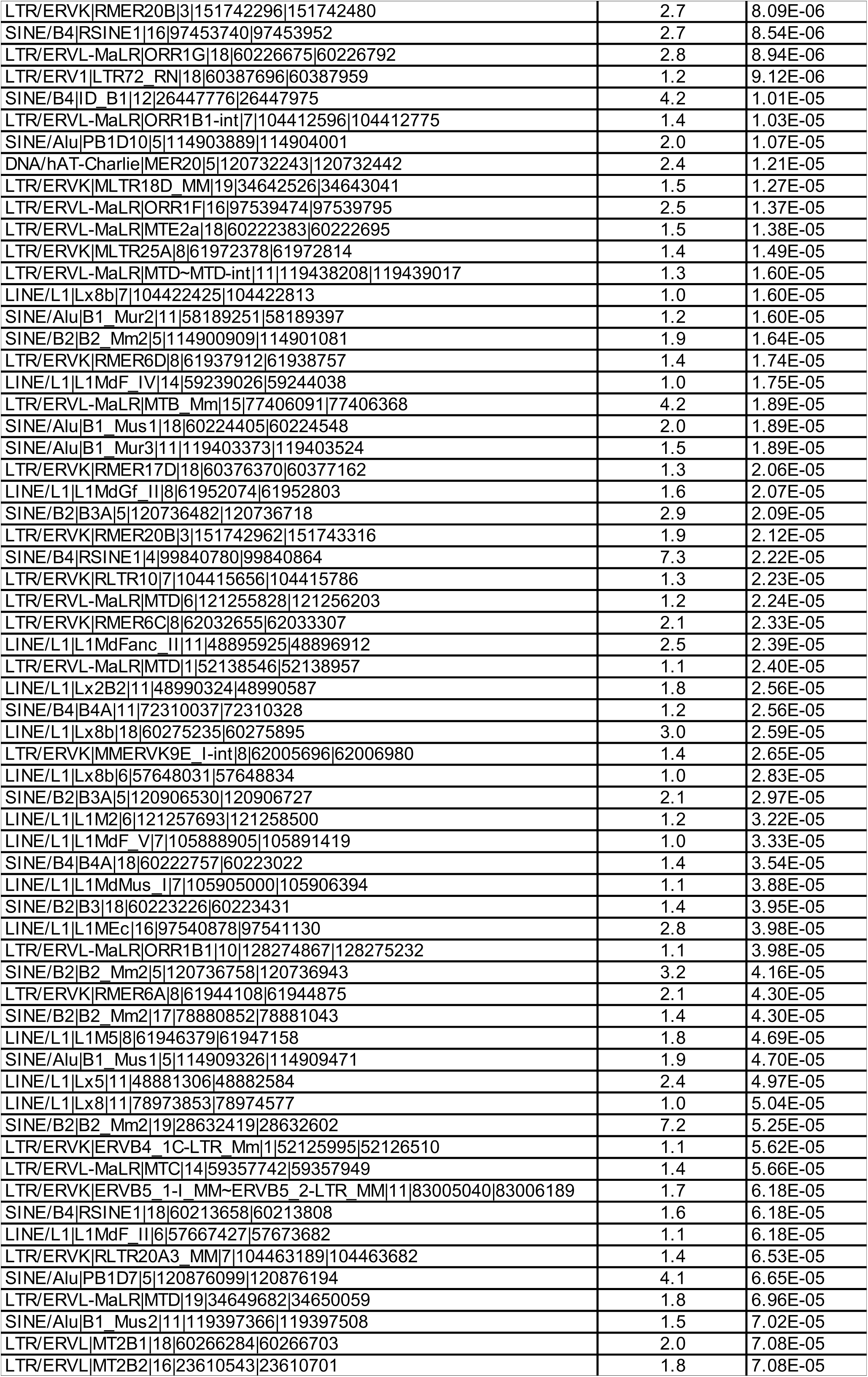
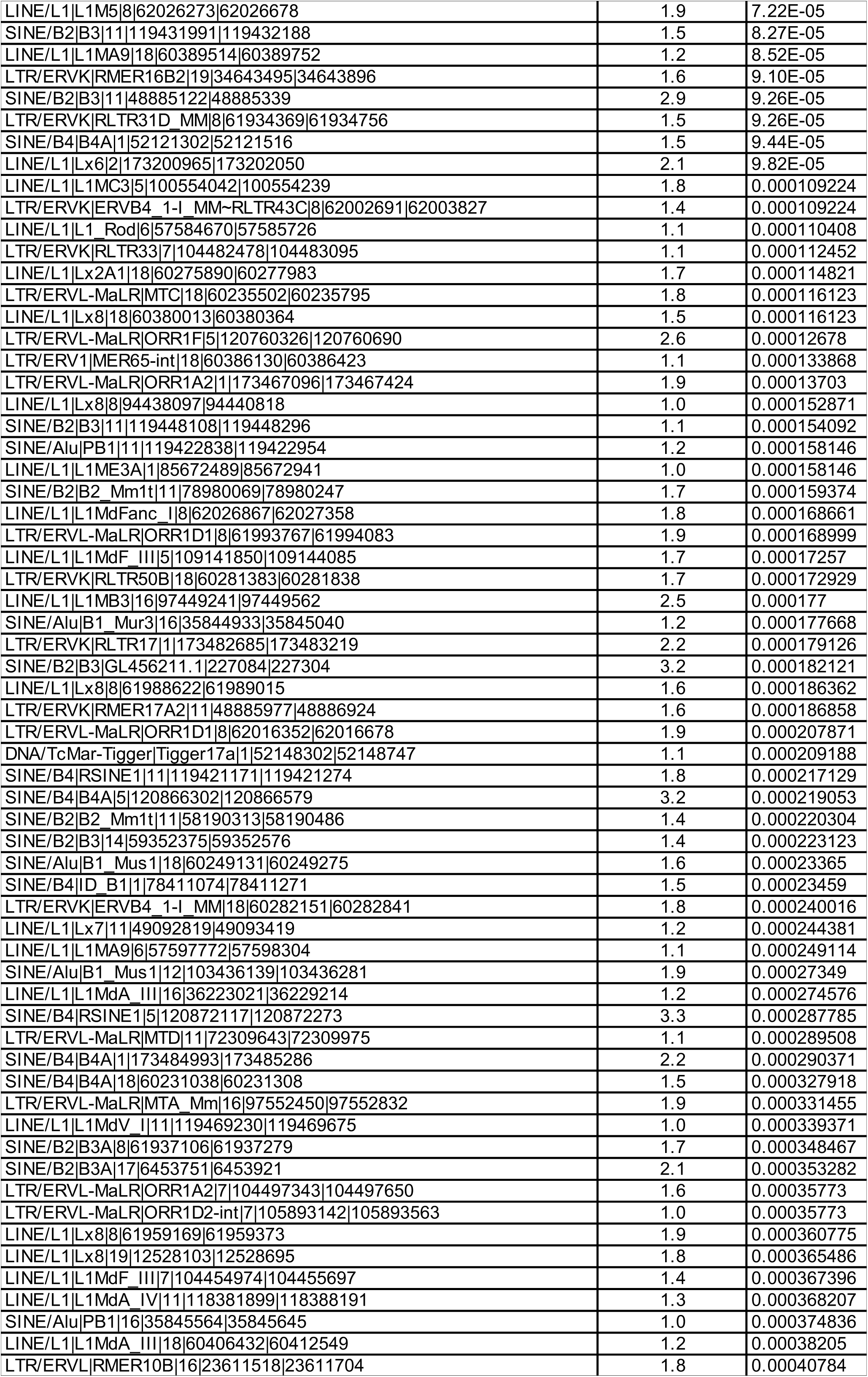
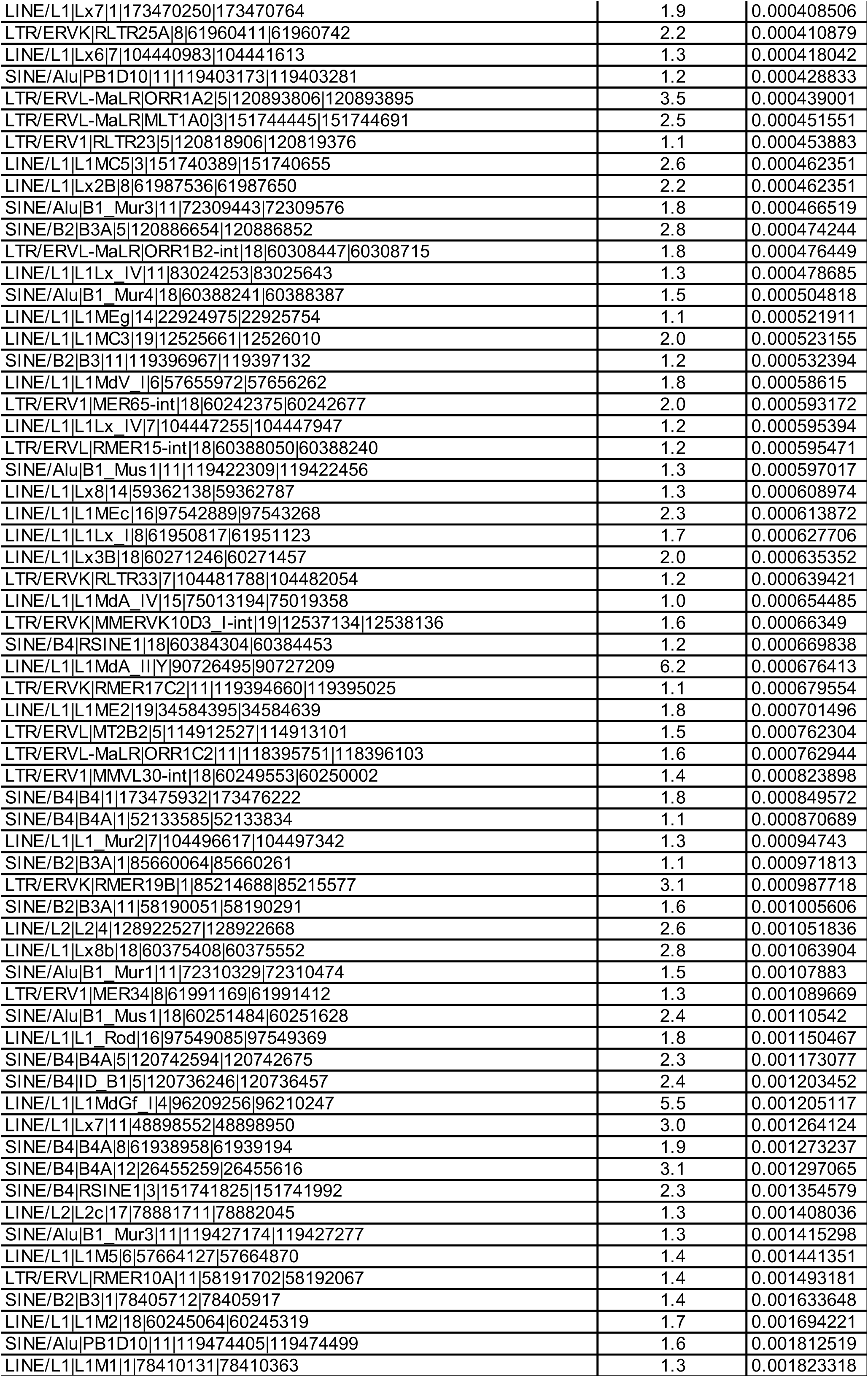
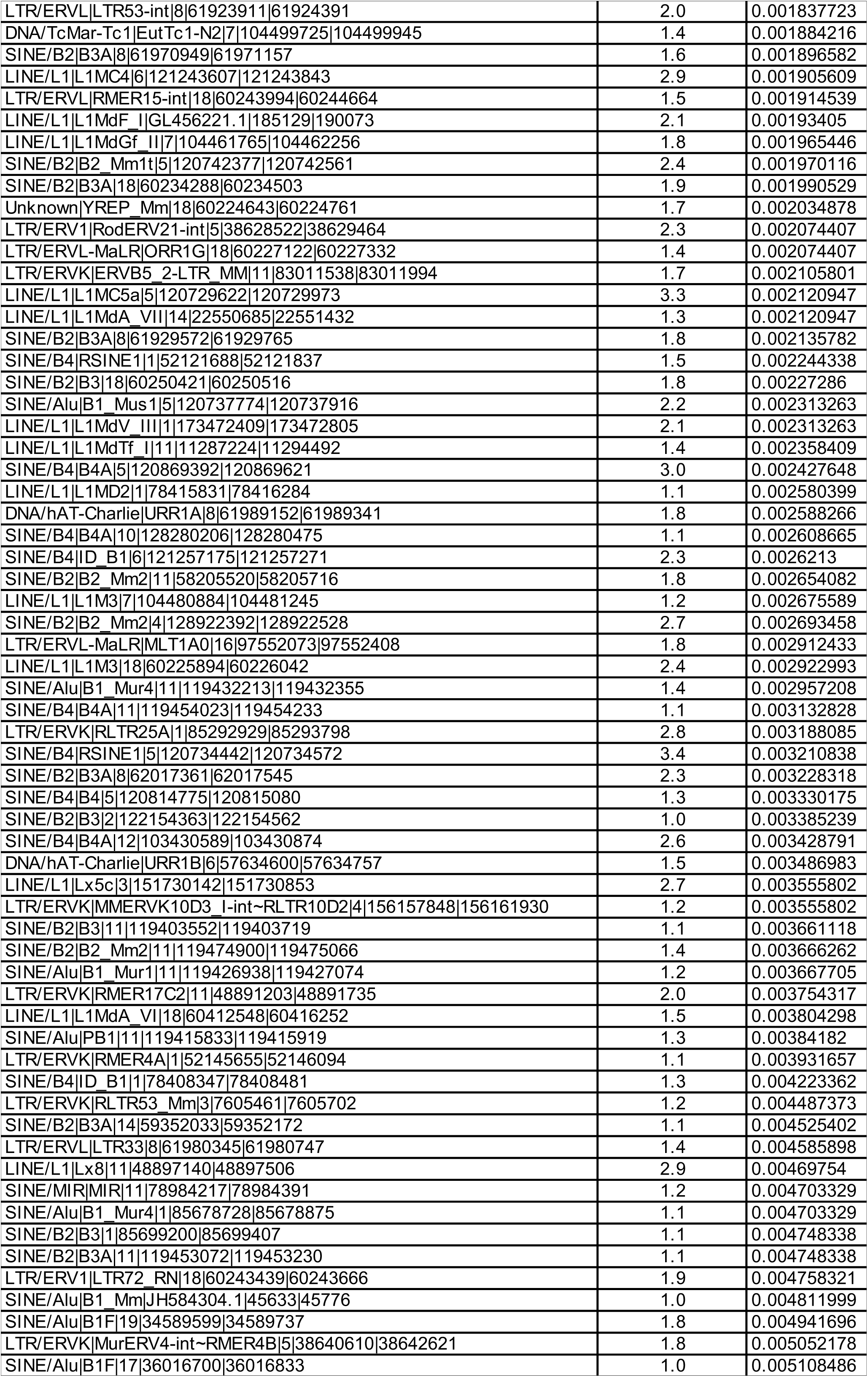
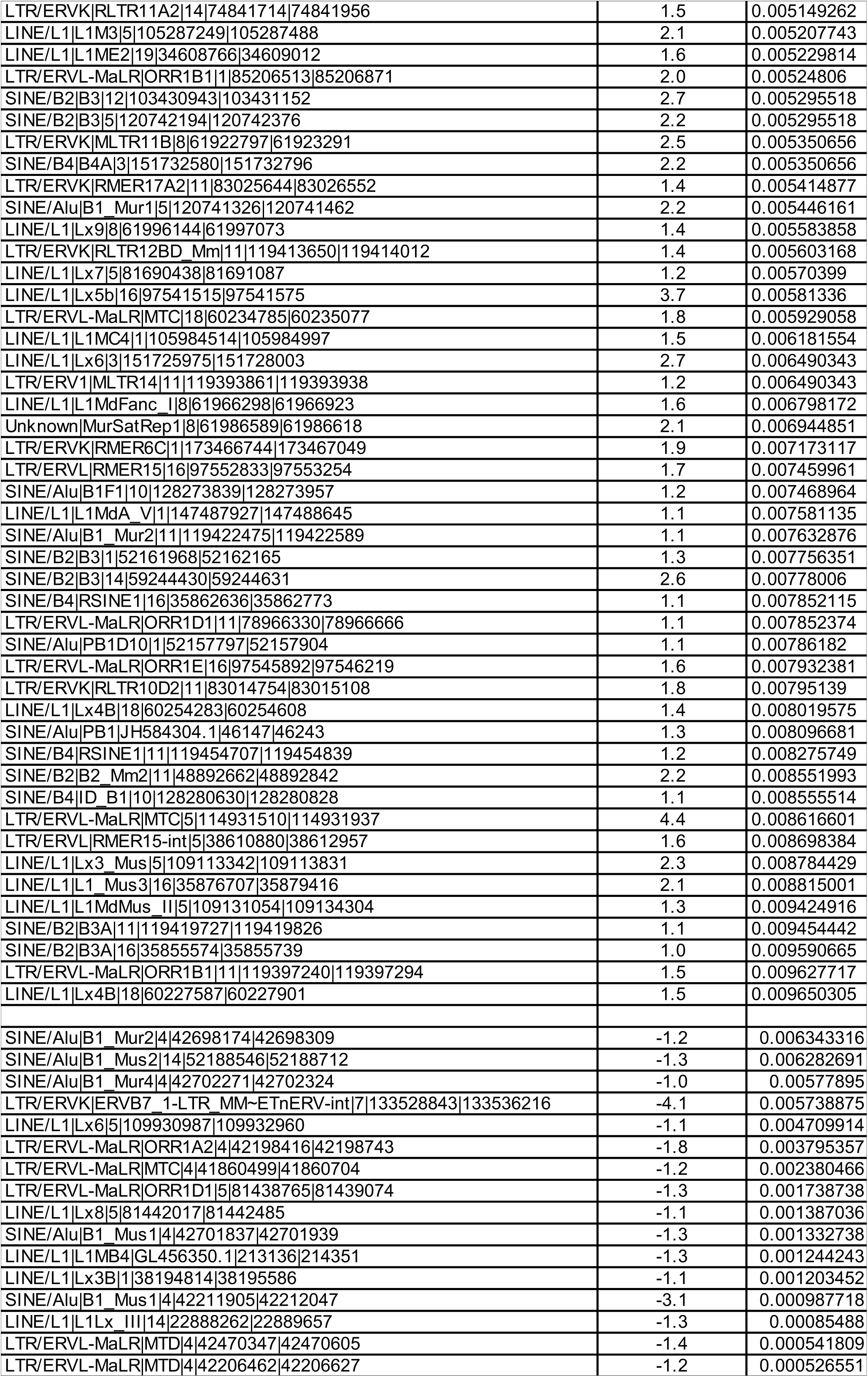
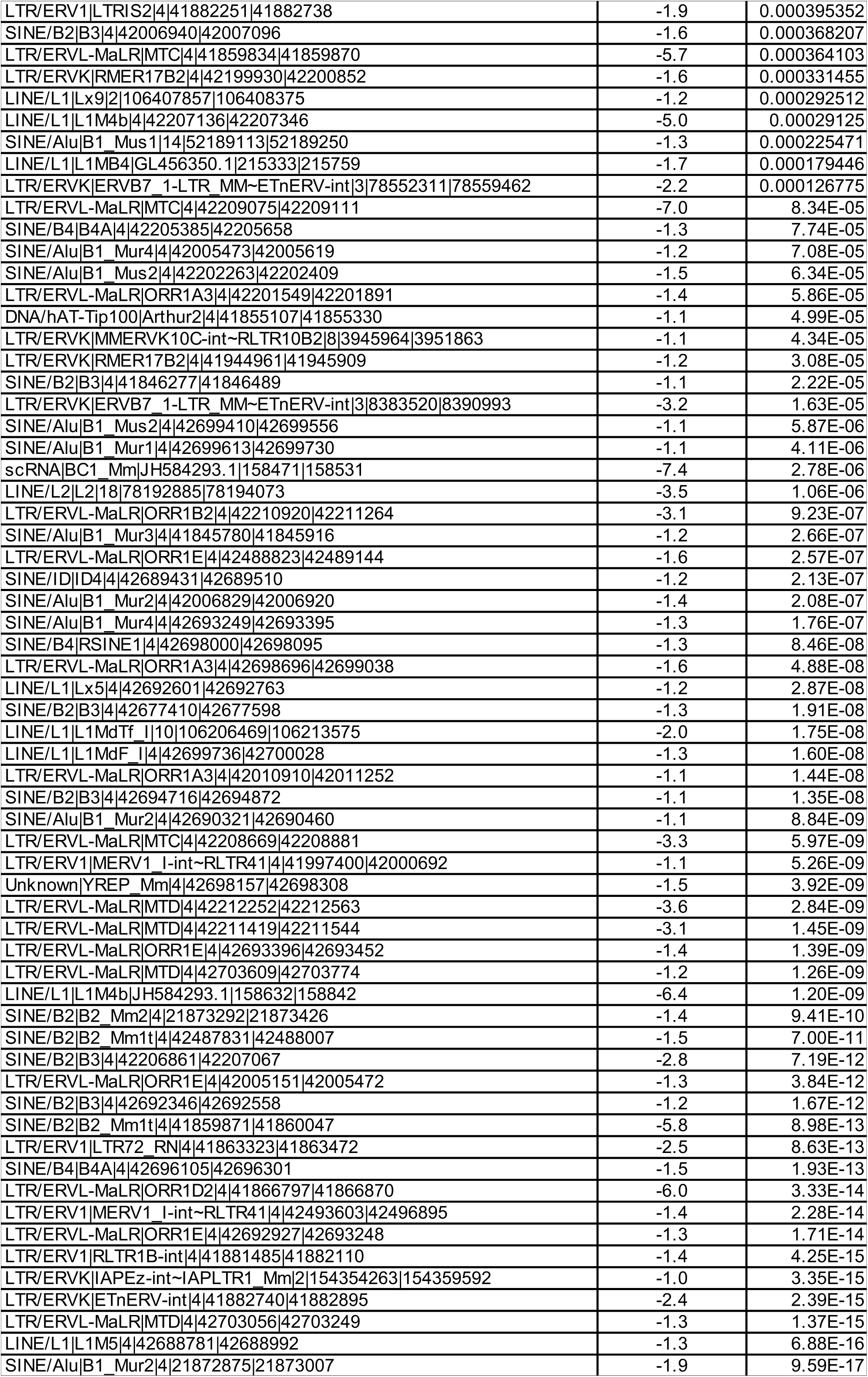
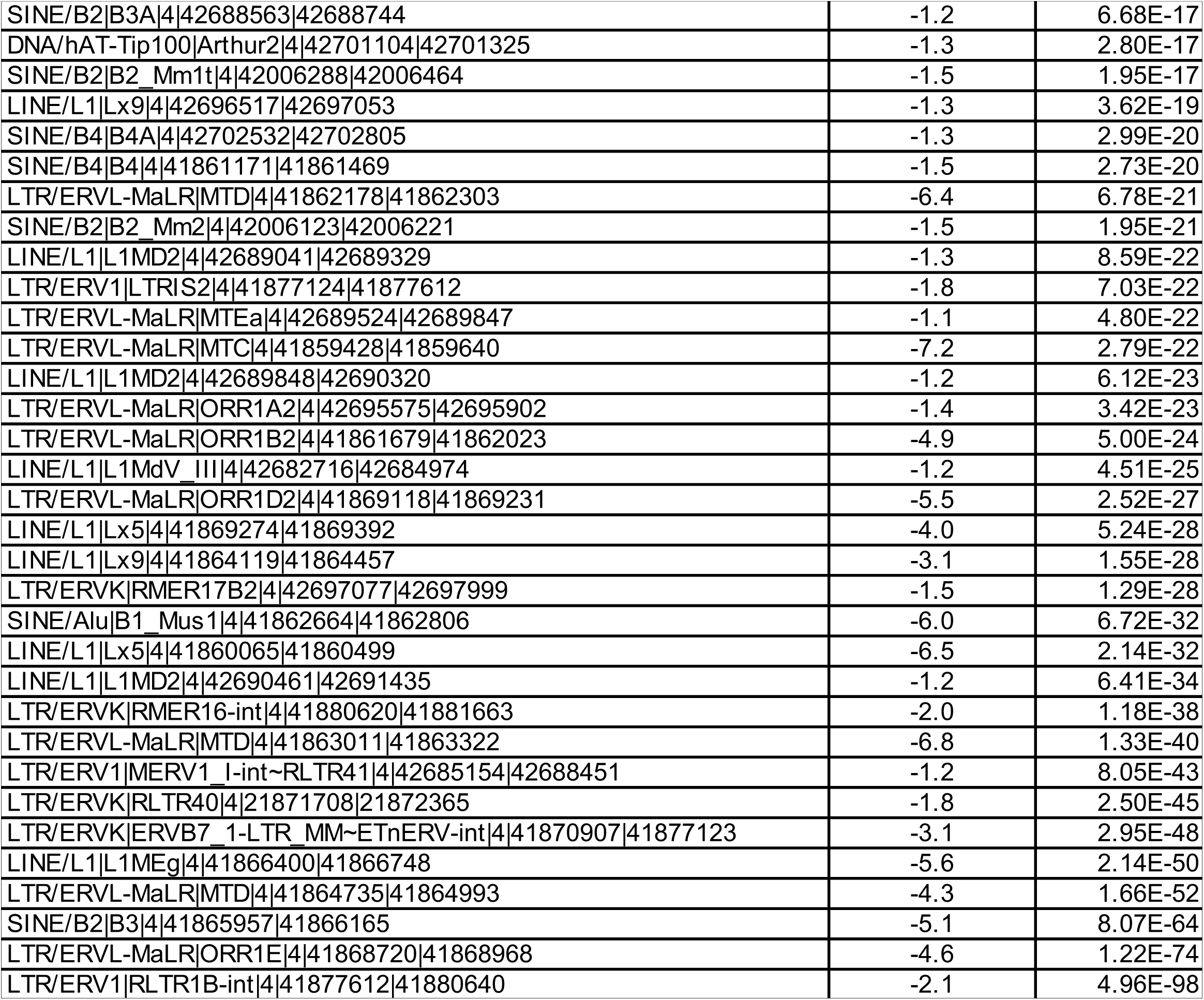
Differentially expressed REs. REs differentially expressed in *Adar1 ^mZ α/mZ α^* lungs are shown with their log2 fold change and adjusted p-value. The RE class, family and chromosomal location are indicated. REs were sorted by adjusted p-value. The cut-offs used were: log2 fold-change >1 or <-1, and adjusted p-value <0.01.

**Table S2.**
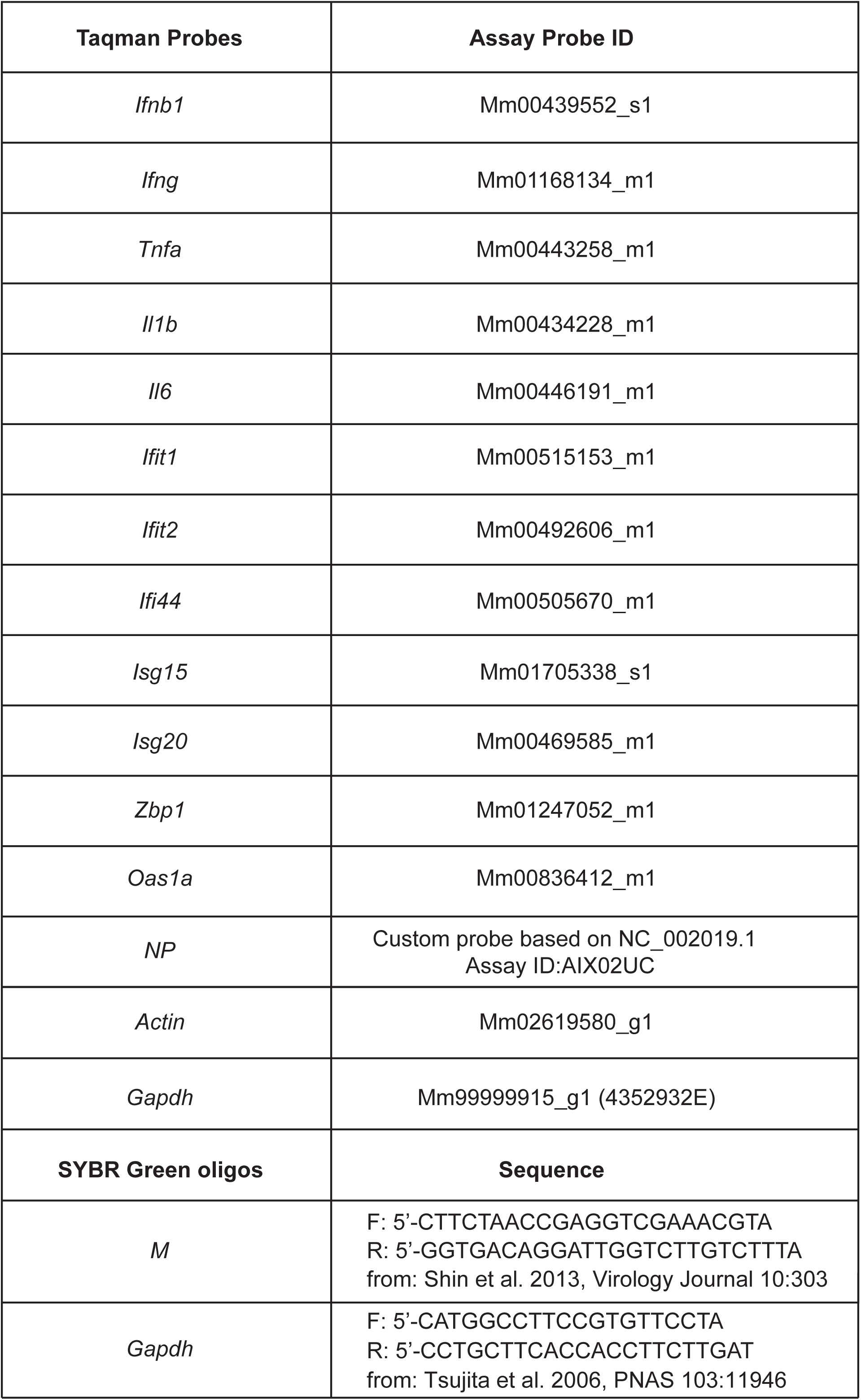
qPCR probes and primers.

